# Single-cell transcriptomics reveal differences between chorionic and basal plate cytotrophoblasts and trophoblast stem cells

**DOI:** 10.1101/2024.07.12.603155

**Authors:** Robert Morey, Francesca Soncin, Sampada Kallol, Nirvay Sah, Zoe Manalo, Tony Bui, Jaroslav Slamecka, Virginia Chu Cheung, Don Pizzo, Daniela F. Requena, Ching-Wen Chang, Omar Farah, Ryan Kittle, Morgan Meads, Mariko Horii, Kathleen Fisch, Mana M. Parast

## Abstract

Cytotrophoblast (CTB) of the early gestation human placenta are bipotent progenitor epithelial cells, which can differentiate into invasive extravillous trophoblast (EVT) and multinucleated syncytiotrophoblast (STB). Trophoblast stem cells (TSC), derived from early first trimester placentae, have also been shown to be bipotential. In this study, we set out to probe the transcriptional diversity of first trimester CTB and compare TSC to various subgroups of CTB. We performed single-cell RNA sequencing on six normal placentae, four from early (6-8 weeks) and two from late (12-14 weeks) first trimester, of which two of the early first trimester cases were separated into basal (maternal) and chorionic (fetal) fractions prior to sequencing. We also sequenced three TSC lines, derived from 6-8 week placentae, to evaluate similarities and differences between primary CTB and TSC. CTB clusters displayed notable distinctions based on gestational age, with early first trimester placentae showing enrichment for specific CTB subtypes, further influenced by origin from the basal or chorionic plate. Differential expression analysis of CTB from basal versus chorionic plate highlighted pathways associated with proliferation, unfolded protein response, and oxidative phosphorylation. We identified trophoblast states representing initial progenitor CTB, precursor STB, precursor and mature EVT, and multiple CTB subtypes. CTB progenitors were enriched in early first trimester placentae, with basal plate cells biased toward EVT, and chorionic plate cells toward STB, precursors. Clustering and trajectory inference analysis indicated that TSC were most like EVT precursor cells, with only a small percentage of TSC on the pre-STB differentiation trajectory. This was confirmed by flow cytometric analysis of 6 different TSC lines, which showed uniform expression of proximal column markers ITGA2 and ITGA5. Additionally, we found that ITGA5+ CTB could be plated in 2D, forming only EVT upon spontaneous differentiation, but failed to form self-renewing organoids; conversely, ITGA5-CTB could not be plated in 2D, but readily formed organoids. Our findings suggest that distinct CTB states exist in different regions of the placenta as early as six weeks gestation and that current TSC lines most closely resemble ITGA5+ CTB, biased toward the EVT lineage.

## Introduction

The development of the human placenta has been a black box, particularly early in gestation, when the organ is difficult to probe, particularly during an ongoing pregnancy(1–3). Following implantation, the cytotrophoblast (CTB) shell expands and invaginations soon give rise to primary and secondary chorionic villi(4, 5). Villous CTB (vCTB) progenitor cells differentiate into two main mature trophoblast types: syncytiotrophoblast (STB), which arise by CTB fusion and serve at the nutrient/gas exchange interface of the floating chorionic villi; and extravillous trophoblast (EVT), which arise through epithelial-mesenchymal transition within the anchoring villi, ultimately invading through decidua and the upper third of the uterine myometrium, remodeling maternal spiral arterioles in order to access oxygenated blood for continuous growth and development of the fetus(6). In 2018, trophoblast stem cells (TSC) were derived from early gestation vCTB, and media formulations developed to allow these cells to either self-renew or differentiate into both EVT and STB(7). This work significantly advanced our ability to study this early “black box” period of placental development, spawning a wealth of publications on factors and pathways that regulate human trophoblast differentiation(8–13). However, while these cells are derived from ITGA6^+^ vCTB, it is unclear how well they represent this bipotential cell population, particularly given their lack of spontaneous differentiation into STB.

Recently, Sheridan et al.(14) compared TSC to trophoblast organoids (T-Org), which can also be derived from early gestation placentas, and are composed of an outer layer of proliferative CTB and an inner layer of spontaneously-formed STB(15, 16). They found that TSC have a distinct transcriptome, expressing several markers of cells in the proximal column of anchoring villi, regions which harbor cells transitioning into EVT(14). Shannon et al. applied single cell RNA-sequencing to compare T-Org and TSC grown in 3D as organoids (TSC-Org) to primary trophoblast within the early gestation placenta and found that TSC-Org show features of both CTB and EVT(17, 18). Here, we set out to evaluate CTB heterogeneity in first trimester human placenta at the single cell level, first comparing early (6-8 weeks) to late (12-14 weeks) first trimester CTB, and subsequently, comparing CTB near the fetal surface (chorionic plate/CP) to those near the maternal surface (basal plate/BP). We identify a bipotential “initial state” CTB which appears to be enriched at the CP, and use a combination of in-situ hybridization and spatial transcriptomics to validate our single cell analysis. We then superimpose single cell transcriptomics from TSC, cultured in 2D, onto our *in vivo* dataset, and determine that these cells represent a rare bipotential progenitor, which resides at the BP and is distinct from the “initial state” CTB. Finally, we characterize TSC using flow cytometry-based analysis of CTB and EVT surface markers, and find that TSC uniformly express proximal column EVT markers ITGA5 and ITGA2, and show that similar cells isolated from first trimester placentas exclusively differentiate into EVT.

## Results

### Transcriptional topography of first trimester trophoblast at single cell resolution revealed heterogeneity of cytotrophoblast based on gestational age

To better understand the transcriptional landscape of first trimester human trophoblast, we performed single cell RNA-seq on four first trimester normal placentas, two early (6-8 weeks gestational age/GA) and two late (12-14 weeks GA). Only the gestational sac and associated villous tissue (no decidual tissues) were dissociated into single cells. In total, approximately 60,000 cells were captured using the 10X Genomics platform and sequenced (**Table S1**). The samples underwent ambient RNA removal, doublet identification and filtering, as well as cell and gene quality control and filtering (See Methods), resulting in approximately 44,000 cells that were then clustered, integrated, and visualized using uniform manifold approximation and projection (UMAP)(19–22) (**Figure S1A**). Leiden clustering resulted in 22 clusters with the non-trophoblast cells generally clustering on the left side of the UMAP and the trophoblast cells generally clustering on the right (**Figure 1A**). Next, marker genes were calculated, and clusters were annotated as either trophoblast or non-trophoblast in origin using the expression of trophoblast specific genes (such as *KRT7, GATA3, PAGE4, TFAP2A, HLA-G, CYP19A1*) and non-trophoblast genes (such as *HLA-A, HLA-B, HLA-DRA, CD14, VIM, CD34*) (**Figure 1B, Table S2**). We were interested exclusively in the trophoblast clusters (0, 2, 3, 4, 5, 8, 10, 11, 13, 14, 17, 19, 21) and removed all non-trophoblast clusters (**Figure S2A**). After the removal of the non-trophoblast clusters, we reclustered the remaining trophoblast cells into 17 clusters and recalculated the marker genes for each cluster (**Figure S2B & S2C, Table S3**).

**Fig. 1.**
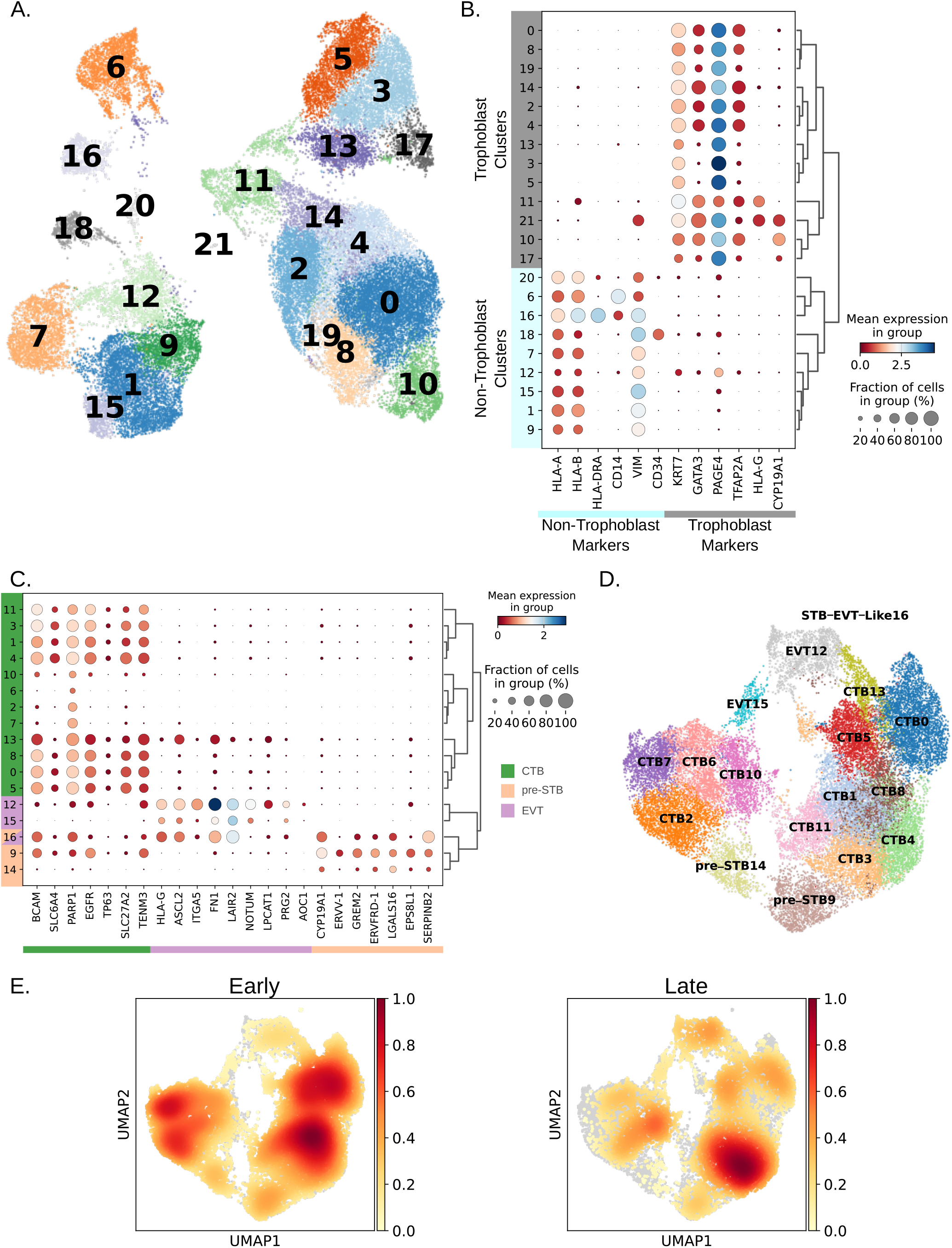
Integration and annotation of early and late first trimester cells. (A) UMAP of all placental cells following quality control filtering, integration, and clustering. (B) Dot plot and hierarchical clustering of clusters using trophoblast and non-trophoblast specific gene expression. (C) Dot plot and hierarchical clustering of clusters using cell type specific gene expression markers. (D) UMAP of trophoblast cells only annotated based on cell type specific gene expression shown in part (C). (E) UMAPs depicting the cell density of the cells in early first trimester (left) and late first trimester (right) placentae.

With our dataset now almost entirely consisting of cells with trophoblast-specific gene expression, we next annotated the clusters using a panel of curated marker genes, into three broad cell types: 1) cytotrophoblast (CTB) (*BCAM, SLC6A4, PARP1, EGFR, TP63, SLC27A2, TENM3*); 2) extravillous trophoblast (EVT) (*HLA-G, ASCL2, ITGA5, FN1, LAIR2, NOTUM, PRG2, AOC1*); and 3) pre-syncytiotrophoblast (pre-STB) (*CYP19A1, ERVV-1, GREM2, ERVFRD-1, LGALS16, EPS8L1, SERPINB2*). We designated cells expressing canonical syncytiotrophoblast (STB) specific genes as pre-STB because STB are large multinucleated cells that presumably would not be able to be captured by the 10X Genomics Chromium platform. This resulted in 17 clusters, including twelve different CTB clusters, two EVT clusters, two pre-STB clusters, and one small cluster (of 15 cells) made up of cells from one, six-week placenta that expressed both EVT and pre-STB marker genes (**Figure 1C, 1D, Table S3**). When compared to all the other trophoblast clusters, the top 200 marker genes from this small cluster were significantly enriched for genes involved in the VEGFA-VEGFR2 signaling pathway and the IL-18 signaling pathway.

Interestingly, the EVT clusters contained more cells from late first trimester placentas while the pre-STB clusters contained more cells from early first trimester placentas. The CTB clusters could be separated into three groups/clades (**Figure 1C**): clade 0, 5, 8, and 13 and clade 2, 6, 7, and 10 were transcriptionally similar, with most of the clusters (with the exception of CTB cluster 10) containing a high proportion of cells from early first trimester placentas. These two clades were transcriptionally distinct from CTB cluster clade 1, 3, 4, and 11, which had a higher proportion of late first trimester cells (**Figure S2D and Table S4**). We also noted that although CTB clusters 2, 6, 7, and 10 did not express any EVT or pre-STB markers, and clustered with the other CTB clusters, only a small fraction of cells expressed pan-cytotrophoblast makers such as *EGFR, TP63*, and *BCAM*, compared to the other CTB clusters (**Figure 1C**).

To better understand the differences between the cells comprising each cluster, we determined the cell density of the early- and late first trimester cells (**Figure 1E and Table S4**). We then determined the differences between the CTB clusters made up of predominantly early or late first trimester cells by comparing CTB8 (early) and CTB3 (late), as well as CTB7 (early) and CTB10 (late) (**Table S5 and Table S6**). In the first comparison (CTB8 vs CTB3), using genes upregulated in the early clusters by at least a log2 fold change of 1.5, we found that the epithelial mesenchymal transition (EMT) and hypoxia pathways were significantly upregulated (adjusted p-value < 0.05). Among the genes upregulated in the EMT pathway were *IGFBP3, CCN1, TPM1, HTRA1, FLNA*, and *MYL9*. In the second comparison, (CTB7 vs CTB10), we found that the MSigDB Hallmark proliferation pathways Myc Targets V1, E2F Targets, and G2-M Checkpoint, were all significantly enriched (adjusted p-value < 0.05) in CTB7 (early)(17). When examining the genes upregulated in the late first trimester CTB clusters, we found that both comparisons significantly upregulated (adjusted p-value < 0.05) the immune-linked inflammatory response pathway, as well as the androgen response and late estrogen response signaling pathways (**Table S7 and S8**)(17).

### Trophoblast trajectory inference identifies a common progenitor CTB cluster

Next, to study the cellular dynamics and relationships between cells and clusters, we performed trajectory inference on our integrated trophoblast cells using RNA velocity(23). In agreement with previous results, our analysis suggested a common CTB initial state (CTB0) with three differentiation trajectories; one to a terminal EVT (EVT12) cell population, one to a pre-STB (pre-STB9) cluster and one to a potentially self-renewing CTB population (CTB10) (**Figure 2A**). Through ranking genes based on their cluster-specific differential velocity expression, being significantly higher or lower compared to the remaining clusters in the population, we identified genes that best explained the RNA velocity vector field. We were interested in marker genes that were differentially expressed in the initial state, which was calculated using CellRank(24, 25) (**Figure 2B**). Genes such as *HMMR, TROAP*, and *LY6E* appeared to be good markers of the initial state/CTB progenitor cells, although *LY6E* expression extended into the EVT12 cluster (**Figure 2C**). Despite integration of our datasets, current implementations of RNA velocity are not designed to yield robust estimates across multiple samples(26). Therefore, we also ran RNA velocity on each patient individually after filtering out all non-trophoblast cells (**Figure S3**), calculating the partial gene likelihoods for each cluster of cells to enable cluster-specific identification of potential drivers, the latent time, which approximates the real time experienced by cells as they differentiate, and the velocity length, which approximates the rate of differentiation (**Figure S4A & B**). We noted that the trajectories from the early placentas had three distinct terminal states, one at the EVT cluster, one at the pre-STB cluster, and one at a CTB cluster marked by the lack of expression of the pan-cytotrophoblast marker *EGFR*. However, it was not clear from trajectories of the two late placentas that the *EGFR*-CTB cluster represented a terminal state or was a source of self-renewing CTB, as seen by velocity vectors (**Figure S3**), the early latent time (**Figure S4A**), and the slower rate of differentiation of this cluster in the late placentas (**Figure S4B**). Moreover, based on the RNA velocity streamlines, it appeared that the early placentae had one well defined bipotential initial state, at cluster 1 in patient 856 and on the border of clusters 5 and 0 in patient 860, whereas in the late placentae, it appeared that there was an EVT progenitor population (cluster 6 in patient 861 and cluster 9 in patient 866) and a distinct pre-STB progenitor population (cluster 0 in patient 861 and cluster 1 in patient 866)(**Figure S3**).

**Fig. 2.**
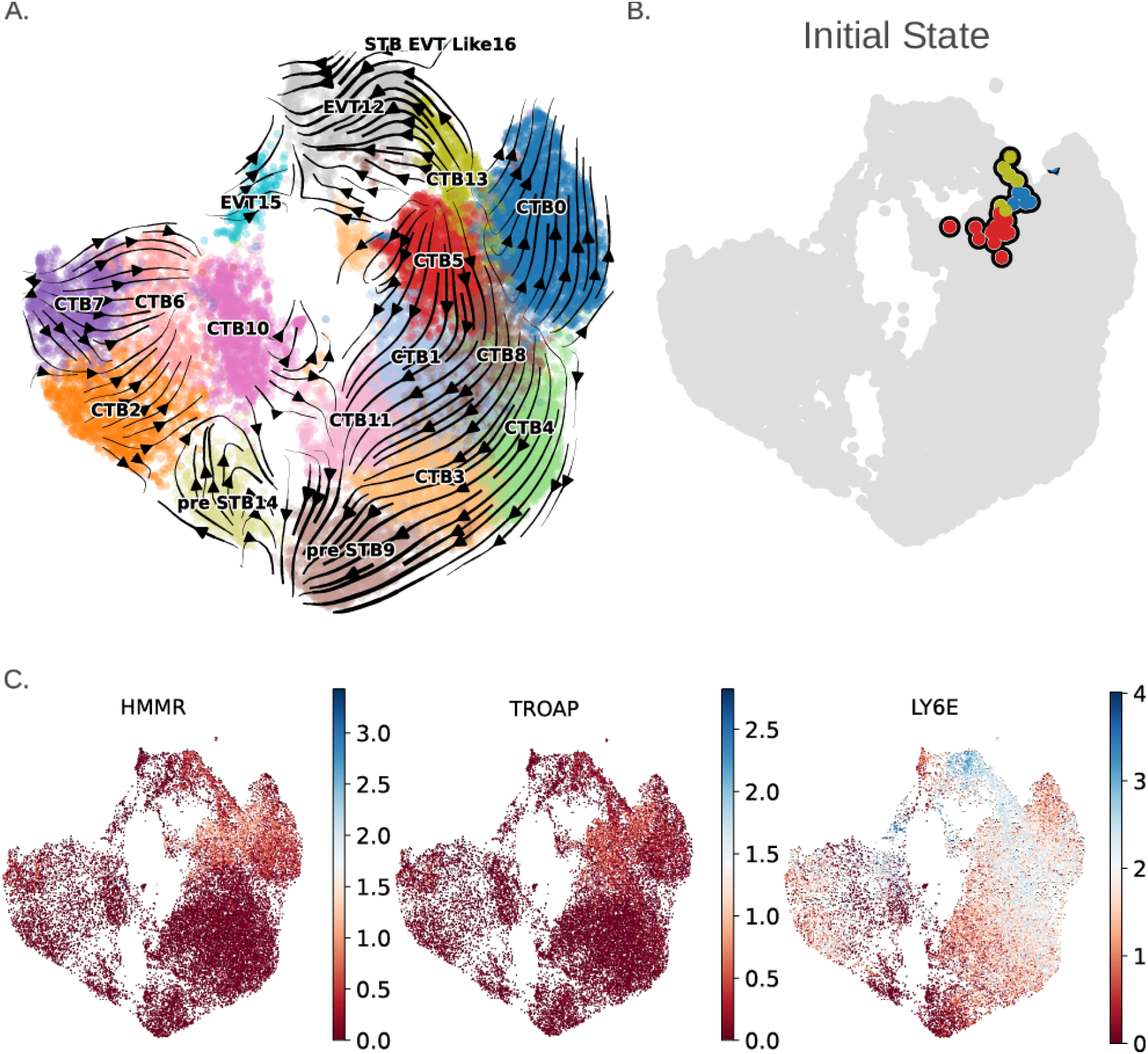
Trajectory inference of first trimester trophoblast cells. (A) RNA velocity projected as streamlines on the integrated first trimester trophoblast UMAP. (B) Initial state as calculated by CellRank. (C) Expression of the initial state markers, *HMMR, TROAP*, and *LY6E* on the integrated first trimester trophoblast UMAP.

To identify cluster-specific potential driver genes for EVT and pre-STB, we looked at the overlap between the four samples of the 100 top-likelihood genes in the clusters that were determined by RNA velocity to be directly preceding either the EVT or pre-STB clusters. For the clusters directly upstream of EVT clusters, there was, not surprisingly, more than double the number of genes in common between patients with placentas of the same gestational age compared to overlap between the early and late placentas. Out of the top 100 genes, only three genes (*LHFPL6, PLS3, TNFAIP3*) were found to be in common between the 4 placenta samples. The overlap between the two early first trimester placentas (19 genes) was enriched for genes related to mitotic cell cycle spindle (adj p-value < 0.05), whereas the overlap between the two late first trimester placentas (24 genes) contained the transcription factors *GATA3* and *RUNX1* (**Table S9**). For clusters directly upstream of the pre-STB clusters, there were only two genes (*TMEM40* and *SEMA6A*) that were found in all four placentas, with the overlap between the two early first trimester placentas (15 genes) consisting of genes enriched for terms relating to extracellular matrix organization (adj p-value < 0.05), and the overlap between the two late first trimester placentas (16 genes) consisting of genes enriched for DNA replication terms (adj p-value < 0.05) (**Table S9**).

### Distinct CTB reside within the basal and chorionic plates

The chorionic (CP) and basal (BP) plate respectively constitute the fetal and maternal surfaces of the placenta, with the chorionic villi emanating from the former and being anchored within the latter. These two surfaces are functionally different but the CTB composition of these two regions has yet to be systematically evaluated. As expected, the initial state of our integrated trophoblast cells was predominantly made up of cells from early first trimester placentas (**Figure 1E and Figure2**). Therefore, to better understand the composition of the cell-types that made up our initial state, we dissected the CP (sac) and BP (anchoring villous tips) regions of two early placentas, isolated CTB using Percoll gradient centrifugation, and performed single cell RNA-seq on these four samples (**Table S1**). Capturing of the two compartments was confirmed by enrichment of *CDX2* and *ASCL2* in the CP and BP cells respectively by qPCR(27, 28). Following integration and removal of non-trophoblast cells (**Figure S5A**), the data were clustered, broadly annotated using the same marker genes as with the whole placenta data, and the composition source of each cluster was calculated (**Figure 3A**). Not surprisingly, the EVT cluster contained a higher percentage of cells from the basal plate and the pre-STB cluster a higher percentage of cells from the chorionic plate. To better understand the differences between the BP and CP CTB, we combined clusters CTB1, CTB2, and CTB7, each of which was almost entirely made up of BP cells and had similar expression profiles, and separately combined the clusters CTB0, CTB3, and CTB8, each of which was almost entirely made up of CP cells and had similar expression profiles (**Figure S5B**). We then performed differential expression analysis using these two groups to find differences between BP and CP CTBs. Interestingly, the genes highly expressed in the basal plate CTB were enriched for the proliferation Hallmark gene set Myc Targets V1 (adj. p-value = 2.2e-10) and Unfolded Protein Response (adj. p-value = 0.0015), whereas the genes highly expressed in the chorionic plate were enriched in the Oxidative Phosphorylation gene set (adj. p-value = 5.7e-20) (**Table S10 and Table S11**). To validate these differences between CP and BP CTB, we used the GeoMx digital spatial profiler, and applied its whole transcriptome atlas (WTA) panel to 3 additional 6-week placentas (**Table S1**), selecting areas of interest (AOI’s) at/near the CP (sac) or BP of these placentas, and segmenting on CTB by EGFR immunostaining (**Figure S5C**). We found that approximately 70% of genes identified as CP- or BP-enriched within our single cell analysis were also differentially expressed between basal and chorionic plate CTB using the GeoMx-based assay (t-test p-value < 0.05) (**Figure S5D**).

**Fig. 3.**
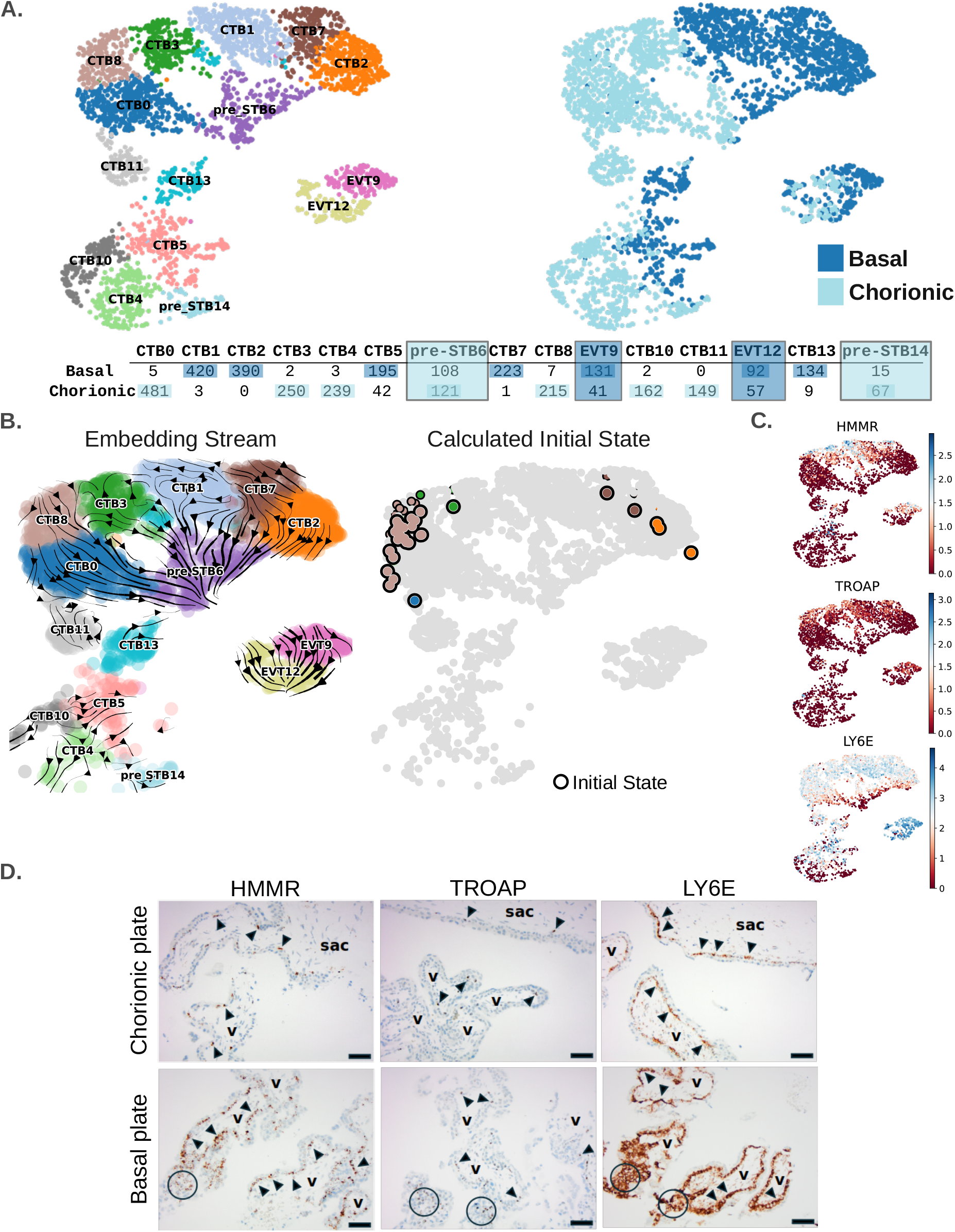
Basal and chorionic plate trajectory inference and initial state. (A) UMAP of annotated and integrated basal and chorionic plate cells from two early first trimester placentae (left). Basal and chorionic plate cells were sequenced from each placenta. When integrated and clustered, basal and chorionic plate cells segregated within each cluster. Location of basal (dark blue) and chorionic (light blue) cells are shown (right). (Bottom) Table shows the number of cells originating from either the basal or chorionic plate making up each cluster. (B) RNA velocity projected as streamlines on the integrated basal and chorionic plate UMAP (left). CellRank calculated initial state locations colored by CTB cluster color (right). (C) Expression of the initial state makers, HMMR, TROAP, and LY6E on the integrated basal and chorionic plate UMAP. (D) In-situ hybridization of the initial state markers, HMMR, TROAP, and LY6E in a 6-week placenta, showing expression in a subgroup of villous CTB (arrowheads) within the gestational sac (“sac”), as well as in chorionic villi (“v”) near the chorionic and basal plates, and within the proximal column trophoblast (circles). Scale bar=125 *µ*m.

To better understand the initial state in our integrated basal and chorionic plate cells, we again performed trajectory inference and calculated the initial state (**Figure 3B**). Both the RNA velocity projected as streamlines on the UMAP, and the CellRank-calculated initial state, indicated that although enriched in the chorionic plate, there were progenitor cell populations in both the basal (CTB cluster 2/7) and chorionic plate CTB (CTB cluster 8), both of which appeared to be bipotential progenitor cell populations that differentiated to both EVT and pre-STB. Additionally, we were interested in how well our initial state markers were expressed in the two basal and chorionic plate progenitor cell populations and found that as in our whole placenta dataset, these genes were highly expressed in what appeared to be our initial state CTB cell populations (**Figure 3C**). We validated these markers using in situ hybridization of five early first trimester placentae, and confirmed expression of all three of these genes in CTB at or near the chorionic plate and in CTB and proximal column trophoblast at the basal plate, with *HMMR* and *TROAP* being less abundant than *LY6E* transcripts (**Figure 3D**). These results suggest that the progenitor CTB populations in both the basal and chorionic plate may have some transcriptional similarity but that microenvironmental factors play a large role in the differences seen in their progeny.

### TSC integration reveals similarities to initial state CTB as well as unique gene expression

In 2018, Okae et al. derived so-called trophoblast stem cells (TSC) from ITGA6^+^ CTB fraction of early first trimester (6-8 week gestational age) placentae, and demonstrated their capacity to differentiate into both EVT and STB(7). Since then, many groups, including ours, have used these or other similarly-derived cell lines, to study various aspects of trophoblast differentiation(29). We therefore sought to compare these TSC to various subgroups of first trimester CTB in order to identify where TSC fall along the developmental pseudotemporal order. We used three TSC lines, derived in our lab from early gestation placentae, and performed scRNA-seq (**Table S1**). Following integration with our placental data, which was pre-filtered for trophoblast (**Figure S6A**), we performed clustering and annotated the clusters based on the same markers we used to annotate the placental trophoblast clusters (**Figure S6B & C, Figure 4A**). Furthermore, in addition to the markers for CTB, EVT, and pre-STB utilized earlier, we examined the expression of the previously identified initial state marker genes in each cluster (**Figure S6C**). Our analysis revealed that clusters TSC0 and TSC12, which were predominantly comprised of cells from TSC lines, exhibited relatively high expression of initial state markers (*TROAP, HMMR*, and *LY6E*), expressed many CTB markers, and in the case of TSC0 (the larger of the two TSC clusters), also expressed many EVT markers. As a result, cluster TSC0 was most similar to our EVT8 cluster and cluster TSC12 was most similar to our CTB10 cluster (**Figure 4A and Figure S6C**). However, marker gene detection revealed striking differences between the two TSC and other trophoblast clusters (**Table S12**). For example, the noncanonical genes *YBX3, CALM1*, and *CALM2* were highly expressed in our TSC clusters relative to our trophoblast clusters, whereas the ubiquitously expressed CTB marker *PAGE4* was not expressed in our TSC clusters (**Figure 4B**). TSC clusters also expressed the proximal column trophoblast/immature EVT markers ITGA2 and ITGA5 (**Figure 4B**). We confirmed by immunostaining that, within first trimester trophoblast, ITGA2 and ITGA5 are confined to anchoring columns (HLA-G^+^), with rare ITGA2^+^ cells identified only within the proximal column, but absent in vCTB and STB (CYP19A1^+^) (**Figure S7A & B**) as previously reported(18, 30). We also performed dual in-situ hybridization for ITGA5 and ITGA2 in first trimester placentae, and confirmed that rare dual-positive cells are only present within the proximal column (**Figure S7C**). Additionally, we performed in-situ hybridization with a PAGE4-specific probe and validated our scRNA-seq data, showing that this gene has the converse expression pattern as ITGA2 and ITGA5, with expression in villous CTB and lacking in trophoblast columns (**Figure 4C**). This suggests that, although our TSC lines showed similarities to villous CTB, they were transcriptionally distinct from their *in vivo* counterparts.

**Fig. 4.**
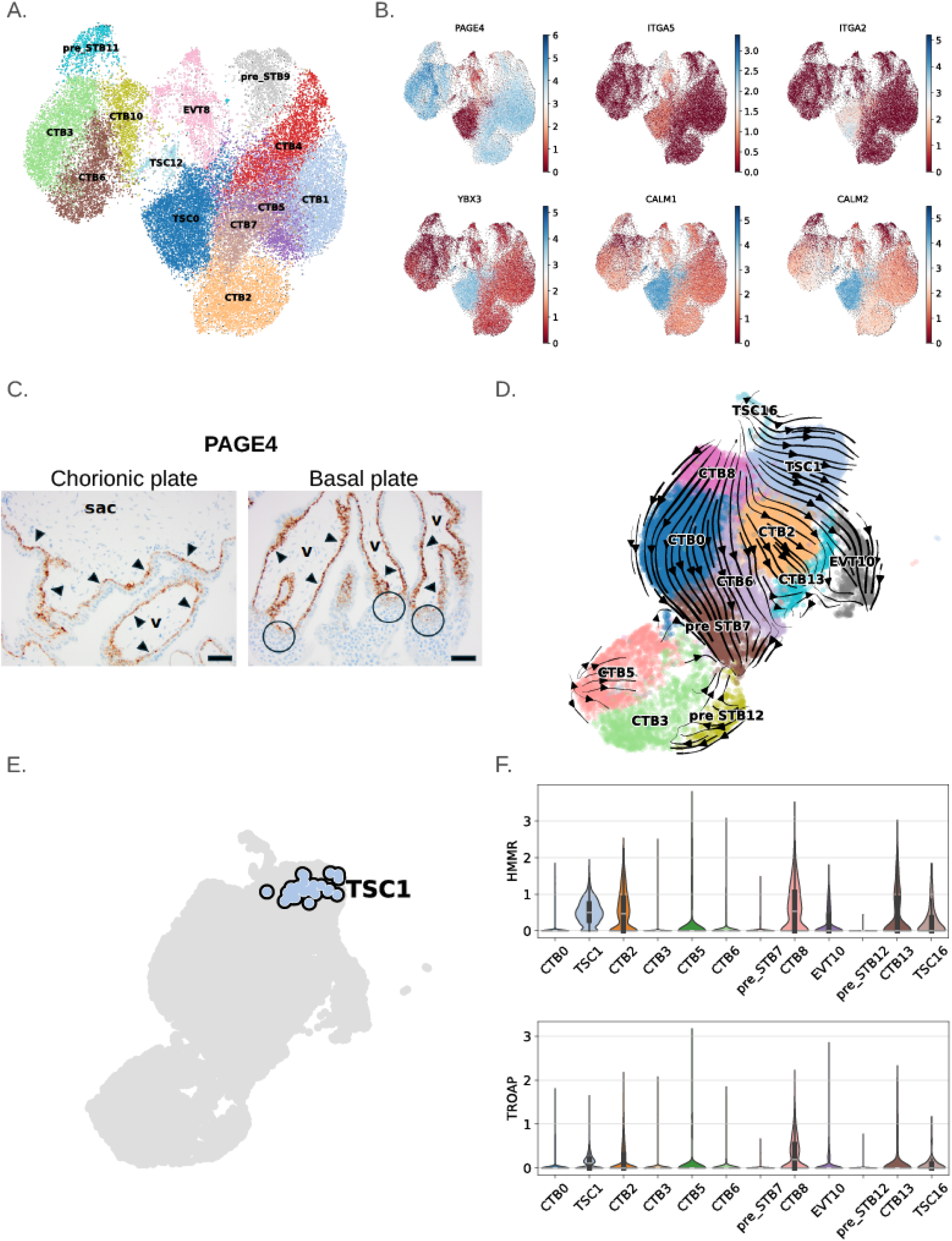
Annotation and trajectory inference of integrated TSC and first trimester trophoblast. (A) UMAP of integrated trophoblast and TSC annotated based on cell type specific gene expression as shown in **Figure S6C**. (B) Differentially expressed genes in TSC cluster compared to placental clusters in early first trimester placentae and TSC integrated UMAPs. (C) In-situ hybridization of *PAGE4* in a 6-week placenta, showing uniform expression in villous CTB (arrowheads) within the gestational sac (“sac”), as well as in chorionic villi (“v”) near the chorionic and basal plates, but excluded from proximal column trophoblast (circles). Scale bar=125 *µ*m. (D) RNA velocity projected as streamlines on the integrated early first trimester placentae and TSC integrated UMAP after the late first trimester placental cells were removed. (E) CellRank calculated initial state location in early placentae and TSC integrated UMAP. (F) Violin/box plots of the initial state genes *HMMR* and *TROAP* in each cluster of the integrated early placentae and TSC data.

We next sought to determine where TSC reside along the trophoblast differentiation trajectory in order to better understand TSC developmental potential. We performed trajectory analysis using RNA velocity with the integrated data but were not able to find a CTB or TSC progenitor population that demonstrated bipotentiality(23) (**Figure S8**). Moreover, the two TSC clusters overwhelmingly integrated with cells from the early (6-8 week), and not the late (12-14 week), first trimester placentas. We therefore combined, preprocessed, filtered for trophoblast cells, and reintegrated the TSC cells with just the early first trimester trophoblast and again performed trajectory analysis using RNA velocity (**Figure 4D**). Using only the integrated TSC and early trophoblast cells, RNA velocity indicated that the TSC clusters and cluster CTB8 represented populations of progenitor cells and Cell-Rank predicted the initial state to be in the TSC1 cluster (**Figure 4E**). We again looked at the same initial state markers (*HMMR, TROAP, LY6E*) in the integrated TSC and early cells and found that both *HMMR* and *TROAP* were specifically expressed in both CTB8 and TSC1 clusters and *LY6E* was highly expressed in these same clusters (**Figure 4F**). Although similar in expression, the CTB8 cells appeared to differentiate into both pre-STB and EVT whereas it was less clear the TSC1 cluster was able to differentiate to pre-STB. To find the differences between the two initial state TSC and CTB clusters, we performed differential expression and found that compared to the *in vivo* CTB8 cluster, the TSC1 cluster was enriched for not only culture-associated terms such as Apical Junction (adj. p-value < 0.00001) but also signaling pathways such as Androgen and Estrogen Response and mTORC1, and the EVT differentiation-related pathway Epithelial Mesenchymal Transition (adj. p-value < 0.00001) (**Table S13**) (**Figure S9A**). The CTB8 cluster was also upregulated for Estrogen Response Late but, in contrast, was also highly enriched in the DNA damage pathway UV Response (adj. p-value < 0.00001), Apoptosis, and the signaling pathways TNF-alpha via NF-kB and KRAS (adj. p-value < 0.001) (**Table S14**) (**Figure S9B**). These results suggest that TSC contain characteristics of progenitor state CTB but have distinct transcriptional programs and may lack the same pre-STB differentiation ability when compared to their villous CTB counterparts.

### TSC express a mixture of CTB and proximal column EVT markers and are distinct from their *in vivo* counterpart in their differentiation potential

To test the growth and differentiation potential of our TSC, we evaluated the expression of both villous CTB (EGFR and ITGA6), proximal column EVT (pcEVT) markers ITGA5 and ITGA2, and pan-EVT marker HLA-G in TSCs. We used six TSC lines derived in our lab from 6-8 week gestation placental tissues (including the 3 subjected to scRNA-seq above), performing flow cytometry for all 5 of these markers, and found these cells to almost uniformly express EGFR (95.5±3%), ITGA6 (94.6±2.4%), ITGA5 (98.6±0.8%), and ITGA2 (99.6±0.4%), with a small fraction co-expressing HLAG (9.4 ±6.2%) (**Figure 5A**). To test whether these markers are retained in TSC following differentiation, we differentiated these cells into EVT or STB, using directed differentiation as described by Okae et al.(7) and performed immunofluorescent staining for ITGA5, along with other markers of differentiation (HLA-G for EVT, and SDC1 and CYP19A1 for STB). We noted that ITGA5 expression was lost following STB differentiation, and retained when TSC were differentiated into EVT (**Figure S10A & B**). These data suggest that, while first-trimester TSCs most closely resemble pcEVT, they do show the expected loss of at least one of these markers (ITGA5) when directed to differentiate into the STB lineage.

**Fig. 5.**
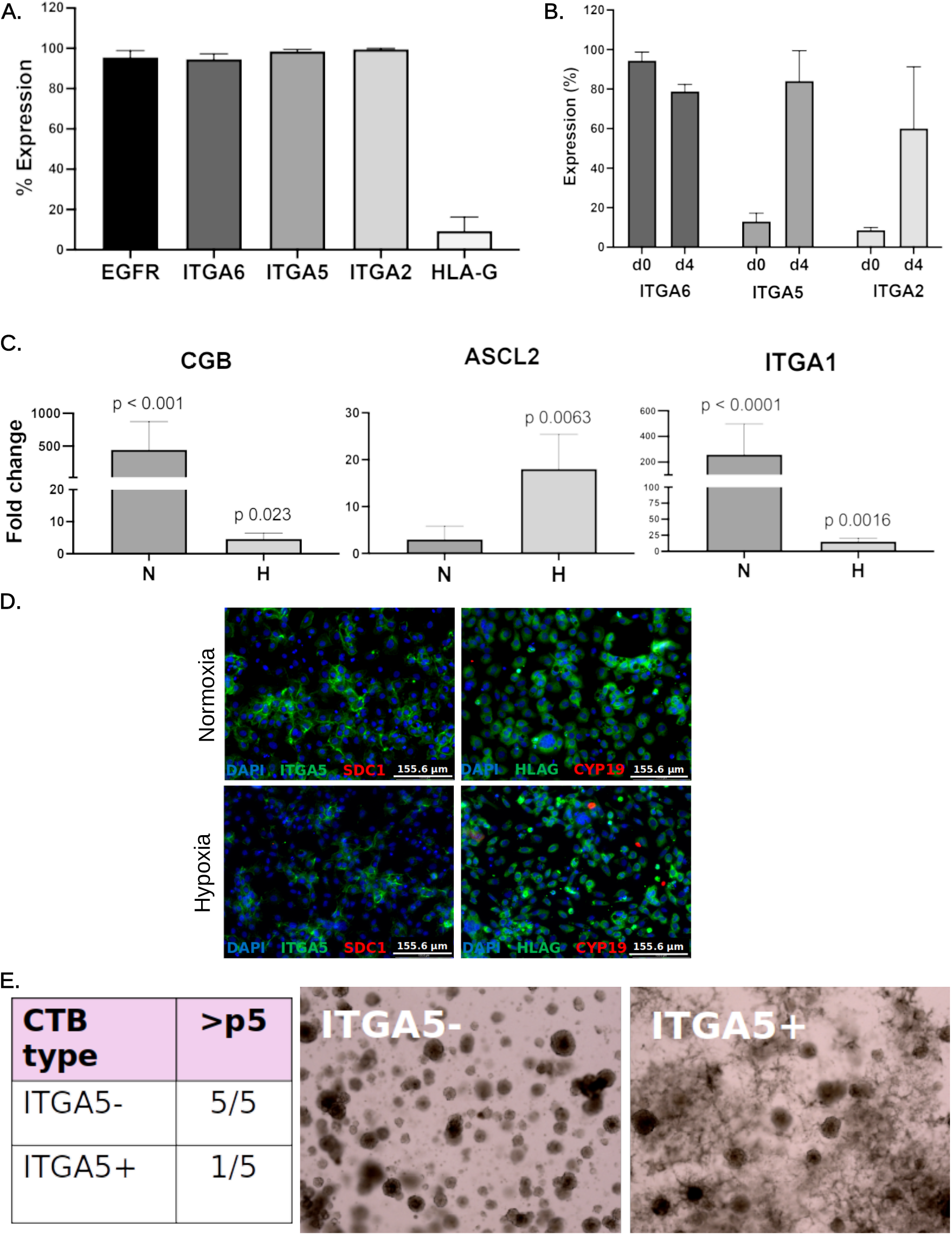
TSC phenotype and differentiation potential compared to primary CTB. (A) Percentage of TSC (undifferentiated state) expressing various markers of CTB and EVT by flow cytometry. Data are reported as mean ± standard deviation of 6 distinct TSC lines. (B) Percentage of first trimester CTB expressing the indicated surface marker by flow cytometry, at isolation and after 4 days of *in vitro* culture. Data are reported as mean ± standard deviation (n=2). (C) Quantitative PCR of STB marker CGB, and EVT markers ASCL2 and ITGA1, in TSC spontaneously differentiated over four days in normoxia (N, 21% oxygen) or hypoxia (H, 2% oxygen). Data are presented as fold change over undifferentiated TSC, reported as mean ± standard deviation (n=2) with p-values calculated using one-way ANOVA. (D) Primary first trimester CTB MACS-sorted based on ITGA5 expression, allowed to spontaneously differentiate over 4 days in either normoxia (21% oxygen) or hypoxia (2% oxygen), then fixed and stained for EVT (HLA-G) and STB (SDC1, CYP19A1) markers, and DAPI. Only ITGA5^+^ cells are shown, as ITGA5^-^ cells did not adhere to any substrate in 2D. (E) Organoid (T-Org) formation using first trimester CTB sorted for ITGA5. Table shows number of successful attempts to derive and culture organoids beyond 5 passages, starting from first trimester CTB MACS-sorted based on ITGA5 expression. Brightfield images of representative T-Org, derived from ITGA5^-^ and ITGA5^+^ CTB.

To determine how culture conditions affected the expression of ITGA5 and ITGA2, we isolated CTB from first-trimester placental tissues and evaluated their integrin profile at isolation and culture after 4 days in 2D. Flow cytometry-based analysis showed a rapid rise of both ITGA5 (d0: 13.05±4.17%; d4: 84.05±15.34%) and ITGA2 (d0: 8.62±1.4%; d4:60.10±31.11%) in just 4 days; expression of ITGA6 was mostly maintained (d0: 94.35±4.31%; d4:78.80±3.54%) during this time period (**Figure 5B**). To better understand the similarities between TSC and the primary CTB most closely resembling TSC, we decided to compare the spontaneous differentiation potential of both cell types. We had previously shown that culture of first trimester CTB, plated on fibronectin in DMEM/F12±FBS media, allowed for differentiation of cells into either STB or EVT over 4 days, depending on oxygen tension (21% vs. 2%, respectively)(31). Therefore, we similarly cultured TSC for 4 days, and found that, by qPCR, they significantly upregulated STB markers, such as CGB, in normoxia and EVT markers, such as ASCL2, in hypoxia (**Figure 5C**); interestingly, however, the EVT marker, ITGA1, was highest when the cells were cultured in normoxia. By morphology and staining, the TSC formed large SDC1^+^ multinucleated cells in normoxia, but preferentially formed HLA-G^+^ mononuclear cells in hypoxia (**Figure S10C**). As a comparison group of CTB, we decided to MACS-sort for ITGA5, since, compared to ITGA2^+^ cells, ITGA5^+^ CTB comprised a larger proportion of isolated mononuclear trophoblast, thus providing sufficient cell numbers from one placenta for one experiment. Only ITGA5^+^ cells adhered to the tested substrates (Col IV or fibronectin), in both oxygen tensions tested (21% or 2%), in either TSC or DMEMF12/FBS media, while very few ITGA5^-^ cells did the same (**Figure 5D**, and data not shown). In both oxygen tensions, after 4 days in culture, ITGA5^+^ cells upregulated the EVT marker, HLA-G, but not the STB markers, SDC1 or CYP19A1, while the few adhered ITGA5^-^ cells did not upregulate any markers (**Figure 5D**, and data not shown). We also tested the ability of CTB, sorted for ITGA5, to form organoids, and found that only ITGA5^-^ organoids showed the proper morphology and could be consistently maintained beyond 5 passages (**Figure 5E**); the majority of the ITGA5^+^ organoids could not be maintained long-term and showed extensive outgrowth formation instead (**Figure 5E**). These data show that ITGA5^+^ TSC are distinct from similar (ITGA5^+^) trophoblast *in vivo*, with the latter lacking bipotentiality, instead able to spontaneously differentiate only into HLA-G^+^ EVT.

## Discussion

Defects in early placental development are thought to be at the root of common pregnancy disorders such as preeclampsia, fetal growth restriction, recurrent miscarriage, and stillbirth. Furthermore, through developmental programming, abnormal early placental development can impact the long-term health of the offspring(32). Despite the critical role early placental development plays in both reproductive success and overall health, our understanding of the development of this transient but consequential organ remains elementary due to ethical constraints, the absence of comparable animal models, and, until recently, the lack of normal (non-transformed) *in vitro* cell models. The derivation of TSC by Okae et al. in 2018 has truly transformed this field, allowing for probing of specific genes and pathways in trophoblast differentiation and function(7, 33– 38). However, recently, this model has come under significant scrutiny, particularly in comparison to the 3D trophoblast organoid (T-Org) models(15, 16), with respect to their differentiation potential(18, 33). Here, we focus on 2D-cultured TSC, derived from 6–8-week gestation placentae, and, using single-cell transcriptomics, compare them to the heterogeneous population of CTB, in both early (6-8 week) and late (12-14 week) first trimester placentae, and within the basal (maternal) vs. chorionic (fetal) plates.

We first compared early (6-8 weeks) to late (12-14 weeks) first trimester CTB and identified three distinct groups of CTB. One set of CTB clusters was comprised primarily of early first trimester CTB, a second of late first trimester CTB, and a third consisted of CTB with very low levels of the canonical CTB marker genes such as *EGFR, BCAM*, and *TP63*, potentially representing an early CTB cell population that has not yet fully matured. When we compared early and late first trimester CTB, we found that the early CTB was enriched in the epithelial mesenchymal transition (EMT) and hypoxia pathways, while the late CTB was enriched for immune-linked inflammatory response pathways. It is well documented that differentiation from CTB to EVT in first trimester placenta involves EMT and hypoxia (hypoxia-inducible factor/HIF-dependent) pathways(31, 39). However, that these pathways are downregulated within CTB with progression of first trimester suggests that, within this short but important time in placental development, the CTB evolve, and likely begin to specialize. In fact, it has been suggested that, while bipotential CTB may exist in early gestation placentae(40), as early as the first trimester, there may be two distinct CTB progenitors, one giving rise to EVT and the other to STB(41–43). Our trajectory analysis of individual placenta samples seems to support these data, showing a definitive bipotential initial state CTB in early first trimester placentae, switching to two initial states in late first trimester tissues, one a precursor of EVT and the other a progenitor of pre-STB. Although further research is needed, including characterization of differentiation potential of such distinct CTB progenitor populations from different gestational age placentae, our data suggest that a prominent bipotential CTB population of cells may exist only in early first trimester placenta and then may either disappear completely or dramatically shrink in number at later gestation.

To our knowledge, this is the first study to directly compare CTB from the chorionic plate (CP) to those in the basal plate (BP) in first trimester placenta by single cell RNA-seq. These two surfaces are known to perform very different tasks in the placenta, but interestingly, have both been suggested to harbor bipotential trophoblast stem cells(30, 44, 45). Not surprisingly, we found significant transcriptional differences between CTB in these two compartments. CP-CTB were enriched for genes involved in oxidative phosphorylation, while BP-CTB exhibited higher expression of proliferation-related gene sets. It was recently reported that CTB differentiation into STB leads to decreased mitochondrial respiration and thus a sharp change in oxidative phosphorylation(46). Interestingly, our trajectory analysis indicated that although CTB in both the CP and BP were identified as “initial state” cells, these cells were enriched in the CP and also appeared to have higher expression of the initial state markers *HMMR* and *TROAP* compared to the CTB in the BP. We verified these findings using both spatial transcriptomics and in situ hybridization. This may indicate that CTB in the BP are primarily a source of EVT precursors, whereas the CTB in the CP are more likely to form STB but still hold the potential to differentiate into EVT. In fact, studies using first trimester villous explant cultures have demonstrated the existence of two distinct CTB subtypes, with CTB at the villous tips (e.g. basal plate) representing a more robust progenitor cells (that is more resistant to cell death *in vitro*), able to give rise to EVT for many days in culture(42).

Recently, several groups have compared TSC to T-Org and *in vivo* trophoblast(8, 14, 18, 47), with a focus on TSC grown in 3D. Similar to our results, these studies have also concluded that the TSC state shares transcriptional hallmarks of proximal column, and not villous, CTB, thus predisposing TSC to differentiate towards EVT, not STB. Interestingly, we found that compared to the initial state early first trimester CTB, our TSC shared significant transcriptional overlap, including expression of the initial state markers *HMMR, TROAP*, and *LY6E* and the canonical villous CTB markers *EGFR* and *ITGA6*. However, we also found large transcriptional differences between these two cell types, with TSC showing upregulation of *YBX3, CALM1*, and *CALM2*, as well as *ITGA2, ITGA5, TAGLN* and *SOX15*, the latter four markers previously highlighted by Shannon et al.(18). Not surprisingly, the genes upregulated in TSC were enriched for terms that are associated with cell culture such as Apical Junction but also for EMT, which as mentioned above, is necessary for EVT differentiation. Our TSC also mirrored EVT in their almost complete lack of expression of the canonical villous CTB marker, *PAGE4*, a stress-response gene with known anti-apoptotic roles in the prostate(48), but otherwise not well-characterized functionally. To confirm our scRNA-seq data, we characterized TSC lines from 6 different placentae, and demonstrated that, in fact, all lines uniformly expressed the pcEVT markers *ITGA2* and *ITGA5*. Notably, our TSC retained the expression *ITGA5*, only upon differentiation to EVT, but lost it with forskolin-induced differentiation into STB. As noted in previous reports, unlike trophoblast organoids, TSC do not spontaneously form STB(14, 18). In fact, when we allowed TSC to spontaneously differentiate (by culture in DMEM/F12+FBS), the cells formed large syncytia that were positive for the STB marker SDC1, but also upregulated ITGA1, an EVT marker in normoxia. ITGA1 has been noted to be increased in human pluripotent stem cell-derived syncytia, considered to be more representative of invasive primitive syncytium(49). A more detailed evaluation of these TSC-derived multinucleated cells is needed, preferably by single nucleus RNA-seq, in comparison to *in vivo* STB, in order to identify their true cell identity.

It is possible that, similar to first trimester explant culture, where villous CTB quickly undergo cell death while basal plate CTB continue to give rise to EVT outgrowths(42), TSC culture conditions favor the stabilization and expansion of an EVT progenitor-like cell type, instead of a more uniformly bipotential vCTB-like cell-type. In fact, in our hands, first trimester CTB cultured in either standard CTB (DMEM/F12+FBS) or TSC media rapidly and significantly increased the expression of both ITGA2 and ITGA5, markers of proximal column/precursor EVT. When sorted based on ITGA5 expression, only ITGA5^+^ CTB adhered, spontaneously differentiating into only EVT-like cells, while ITGA5^-^ cells, could only be cultured in 3D, as T-Org. These data further support the hypotheses put forth above, that TSC are not only biased toward the EVT lineage, but given the lack of ability to spontaneously form bona fide STB, may therefore not be an appropriate model for true STB differentiation. Instead, T-Org consist of cells that better model villous CTB and spontaneously form an inner STB layer(14, 18), though, again, further comparison of multinucleated cells, derived from TSC (cultured in 2D or 3D) vs. T-Org, to *in vivo* trophoblast, using single nucleus RNA-seq, is needed in order to optimally assess STB formation across these models.

In conclusion, our findings offer an in-depth transcriptomic perspective of the first trimester placenta and have significant implications for understanding early developmental stages of this organ, as well as the utility of TSC as a model system. The identification of distinct CTB populations, especially those located at the basal and chorionic plate, and their developmental trajectories provide a valuable resource for future studies of distinct subtypes of first trimester CTB. The integration of TSC data, with its similarities and differences with *in vivo* CTB, highlights the potential and limitations of TSC in recapitulating early trophoblast differentiation. Finally, validation of scRNA-seq data, not just with spatial transcriptomics and in-situ hybridization, but also with primary cell isolation and culture, offers further insights into the actual differentiation potential and functions of distinct CTB progenitor states. Future studies should aim to further elucidate the factors influencing TSC culture and differentiation and develop conditions for culture and characterization of other CTB progenitors within early gestation human placentae.

### Limitations of the study

The 10X Chromium platform used in this study is limited in the size of cell it is able to capture. While differential recovery is not considered an issue for cells below 30 *µ*m in this system, this has not been tested for cells above this size. This most likely excluded capture of mature multinucleated STB or trophoblast giant cells. Moreover, our dataset was compromised of cells from a relatively small number of presumably normal placentas, sequenced at a relatively shallow read-depth compared to bulk RNA-seq, potentially constraining our ability to find important but lowly-expressed genes. Furthermore, trajectory analysis using RNA velocity relies on static snapshots of cellular states at the moment of measurement, and is therefore reliant on only a small number of genes which appear to obey the simple interpretable kinetics used by RNA velocity(26). Lastly, our study was mainly focused on 2D TSC culture and differentiation and did not include in-depth analysis of 3D TSC or trophoblast organoids. Future studies should use single nucleus RNA-seq and compare multinucleated cells within normal tissue, as well as 2D- and 3D-based models of trophoblast, in order to identify the optimal ways to model these mature cell types.

## Acknowledgements

The authors are grateful to all patients who donated tissues for this research. This work was funded by the National Institutes of Health (NIH) (R01-HD089537, R01-HD104805, and R01-HD111787 to MMP; R00-HD091452 to MH; R01-HD096260 to FS) and NIH/NCATS 2UL1TR001442-08 (CTSA).

MH was also supported by UCSD academic senate RG106839. This publication includes data generated at the UC San Diego IGM Genomics Center utilizing an Illumina NovaSeq 6000 that was purchased with funding from a National Institutes of Health SIG grant (S10 OD026929)” as well as “Computational analysis was performed on the Extreme Science and Engineering Discovery Environment (XSEDE) Expanse at SDSC, which is supported by National Science Foundation grant number ACI-1548562 (allocation ID: BIO220095).

## Author contributions

M.M.P., F.S., and R.M. conceived the project. M.M. consented patients and collected tissues. F.S., S.K., N.S., Z.M., T.B., J.S., V.C.C., D.P., D.F.R., C-W.C., O.F., R.K., and M.H. performed the wet lab experiments and associated analysis. R.M., F.S., K.F., and M.M.P. performed the bioinformatic analysis. R.M., F.S., S.K., and M.M.P. wrote the manuscript with feedback from all authors. M.M.P. provided funding for the project.

## Supplementary Figures Supplementary Tables

Table S1: Sample and sequencing metrics.

Table S2: Top 200 marker genes in each cluster in the primary placenta samples.

Table S3: Marker genes after removal of non-trophoblast clusters, reclustering, and annotation of clusters.

Table S4: Cell numbers of each cluster by gestational age for UMAP shown in **Figure 1D**.

Table S5: Genes upregulated (> 1.5 Log2 fold change) in CTB8 (early) compared to CTB3 (late).

Table S6: Genes upregulated (> 1.5 Log2 fold change) in CTB7 (early) compared to CTB10 (late).

**Fig. S1.**
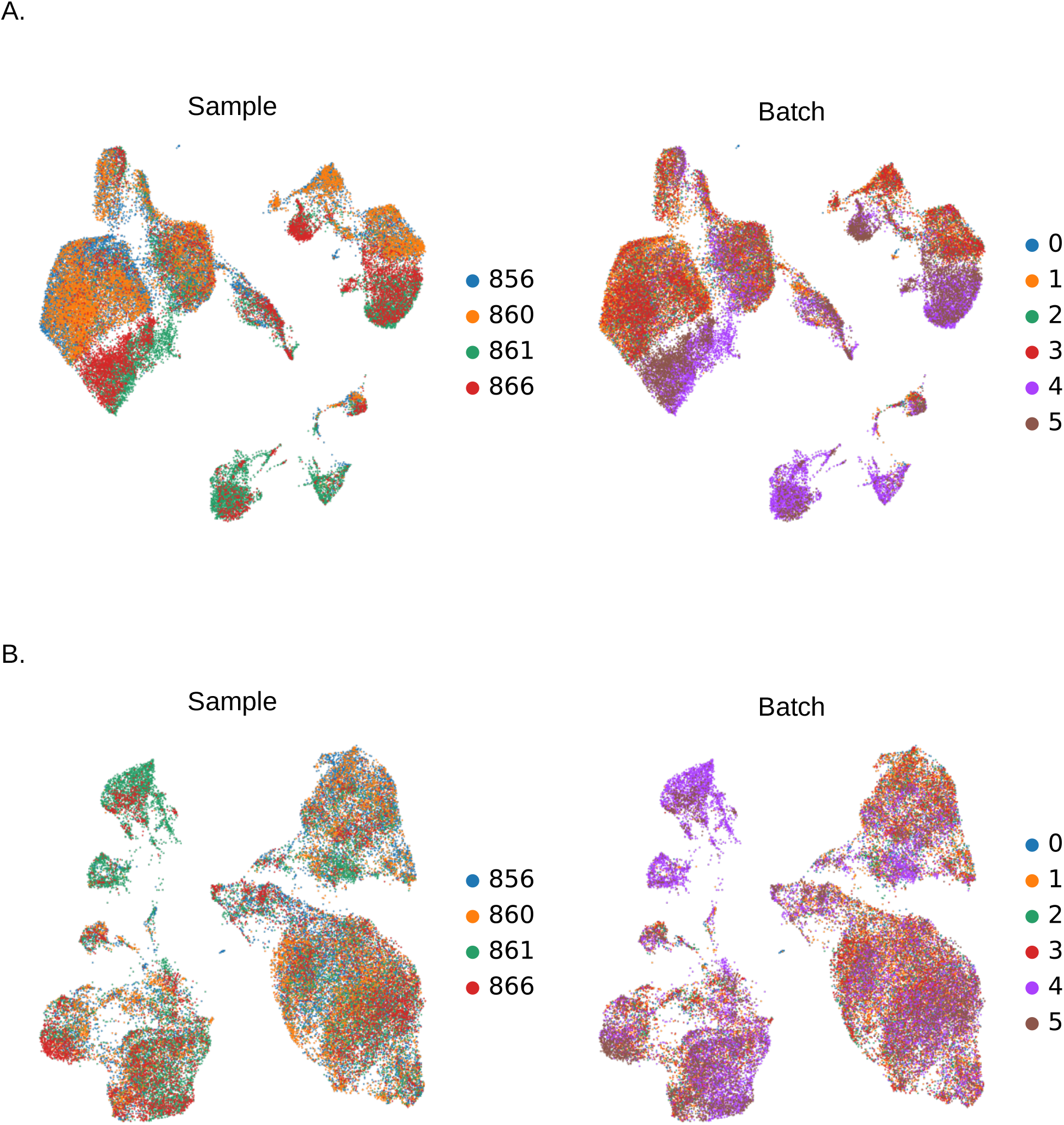
Integration UMAPs. (A) UMAPs of placental samples before integration colored by the patient number (left) and batch number (right). (B) UMAPs of placental samples after integration colored by the patient number (left) and batch number (right)

**Fig. S2.**
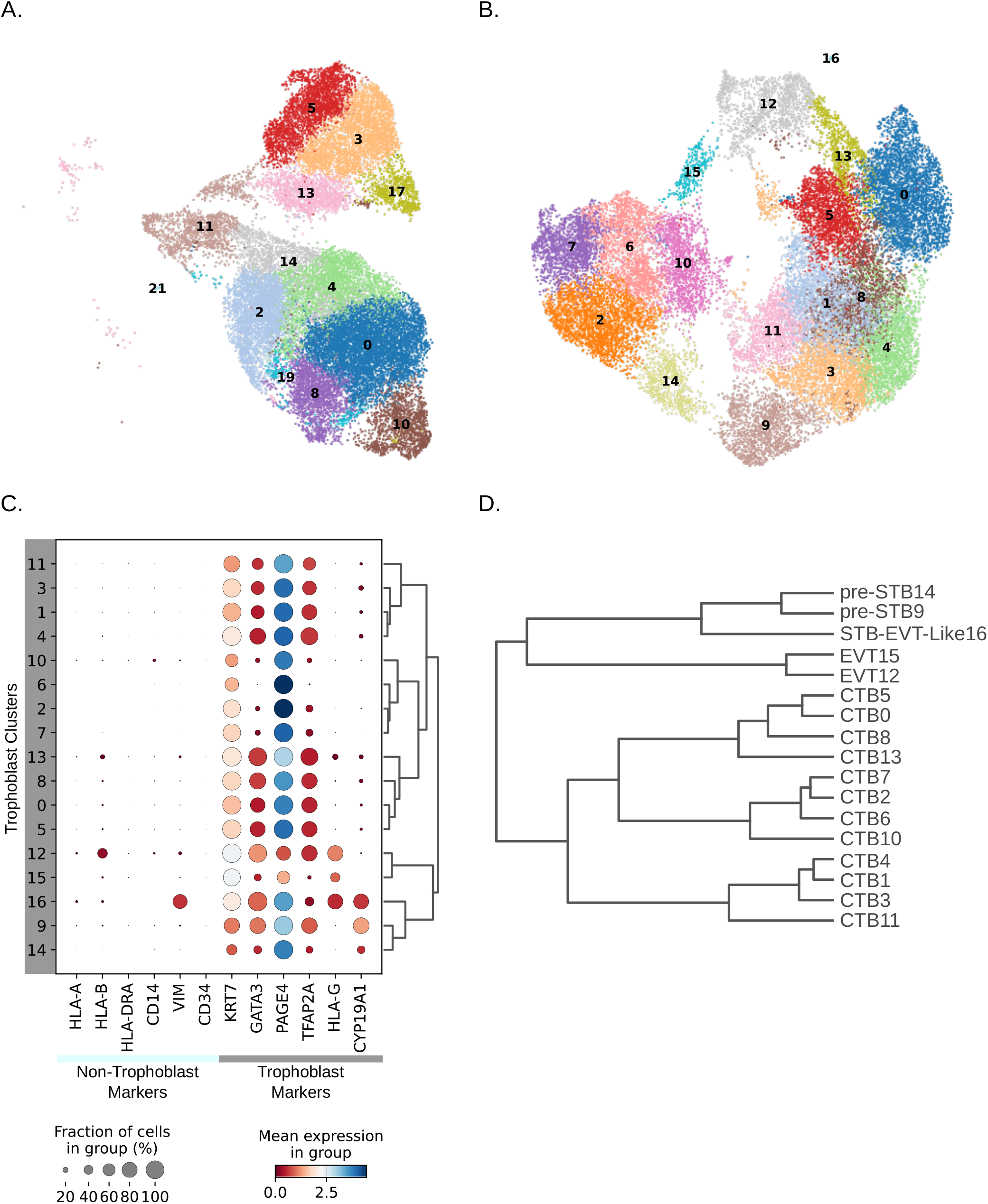
Removal of non-trophoblast clusters and reclustering of trophoblast cells. (A) UMAP following the removal of clusters in Figure 1A deemed to be non-trophoblast cells. (B) Reclustering of cells after removal of most of the non-trophoblast cells. (C) Dot plot and hierarchical clustering showing expression of trophoblast and non-trophoblast specific genes in reclustered UMAP shown in (B). (D) Dendrogram showing the transcriptional similarity between clusters in UMAP shown in (B).

**Fig. S3.**
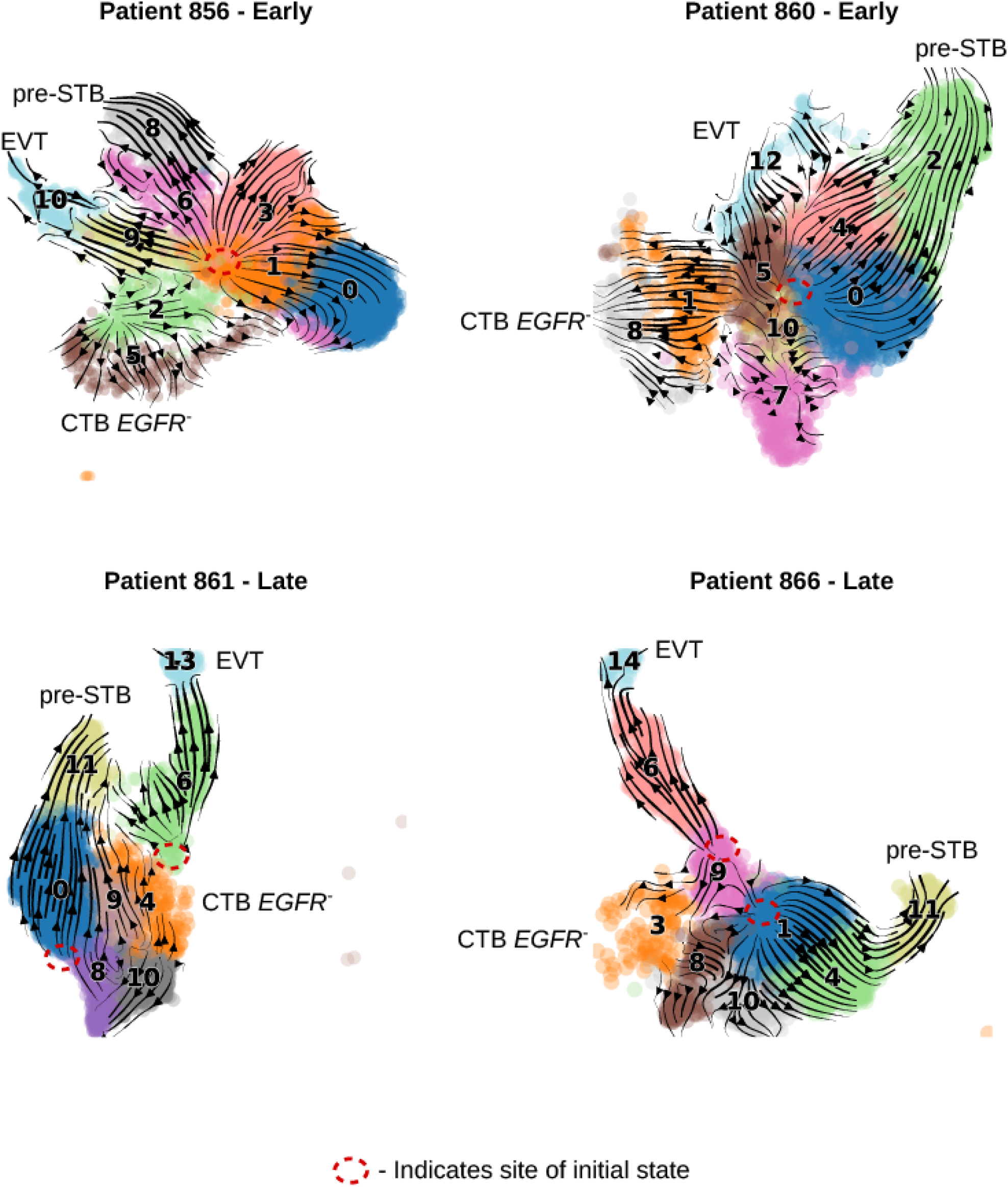
Patient-specific RNA velocity UMAPs. Dynamical RNA velocity projected as streamlines on UMAP embeddings of each of the four individual’s trophoblast cells. The three terminal states in the two early placentas, EVT (HLA-G^+^), pre-STB (CYP19A1^+^) and CTB terminal state (EGFR^-^) were annotated based on the expression of the noted gene(s). In the two late first trimester placentas, the (EGFR^-^) CTB clusters are annotated for comparison purposes.

**Fig. S4.**
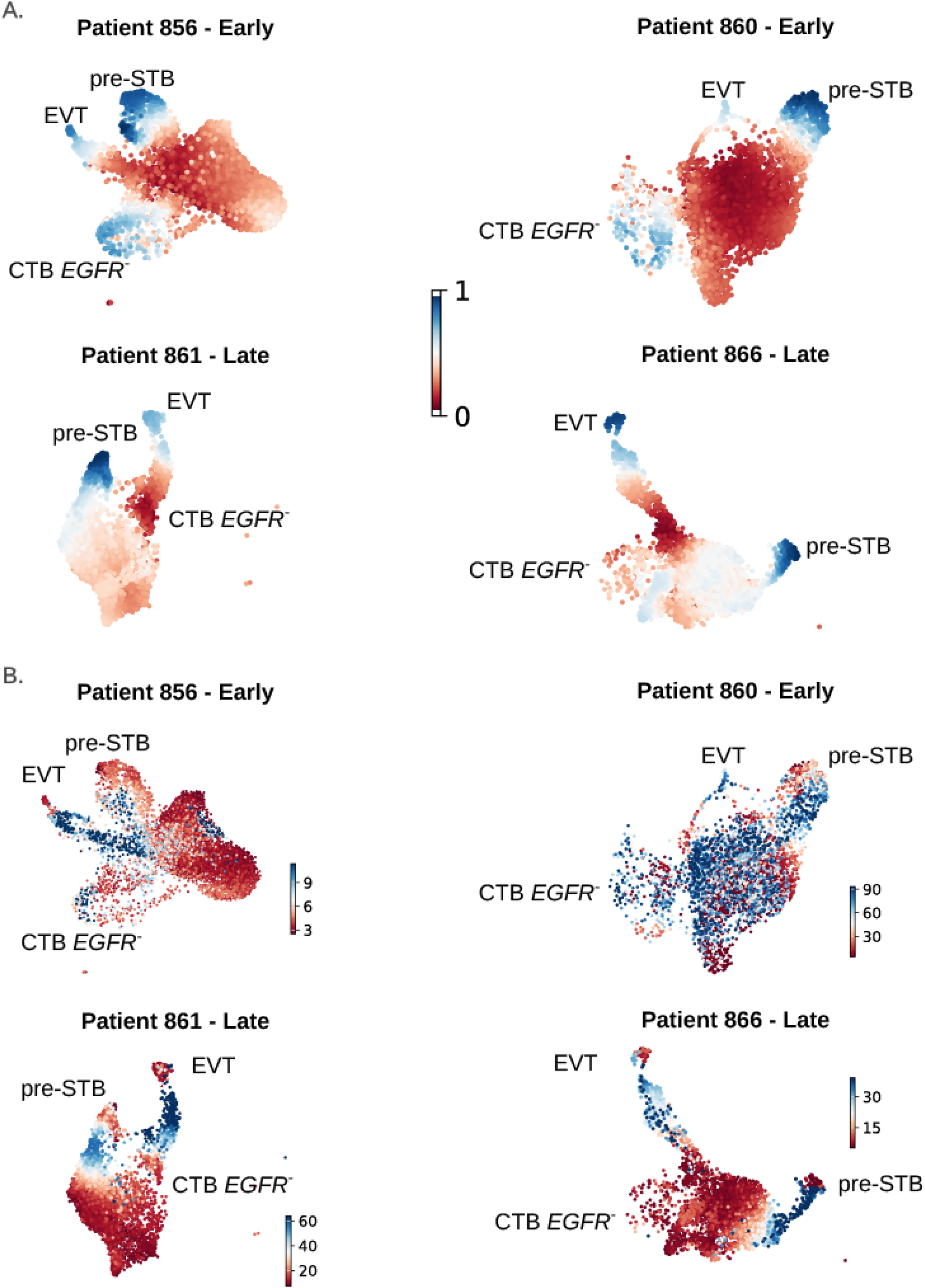
Patient-specific latent time and rate of differentiation UMAPs. (A) UMAPs colored by the latent time of each of the four patient’s trophoblast cells. The latent time represents the cell’s internal clock and approximates the real time experienced by cells as they differentiate, based only on their transcriptional dynamics. (B) UMAPs colored by the rate of differentiation, which is given by the length of the velocity vector. The blue color represents cells that are differentiating faster than the cells marked in red.

**Fig. S5.**
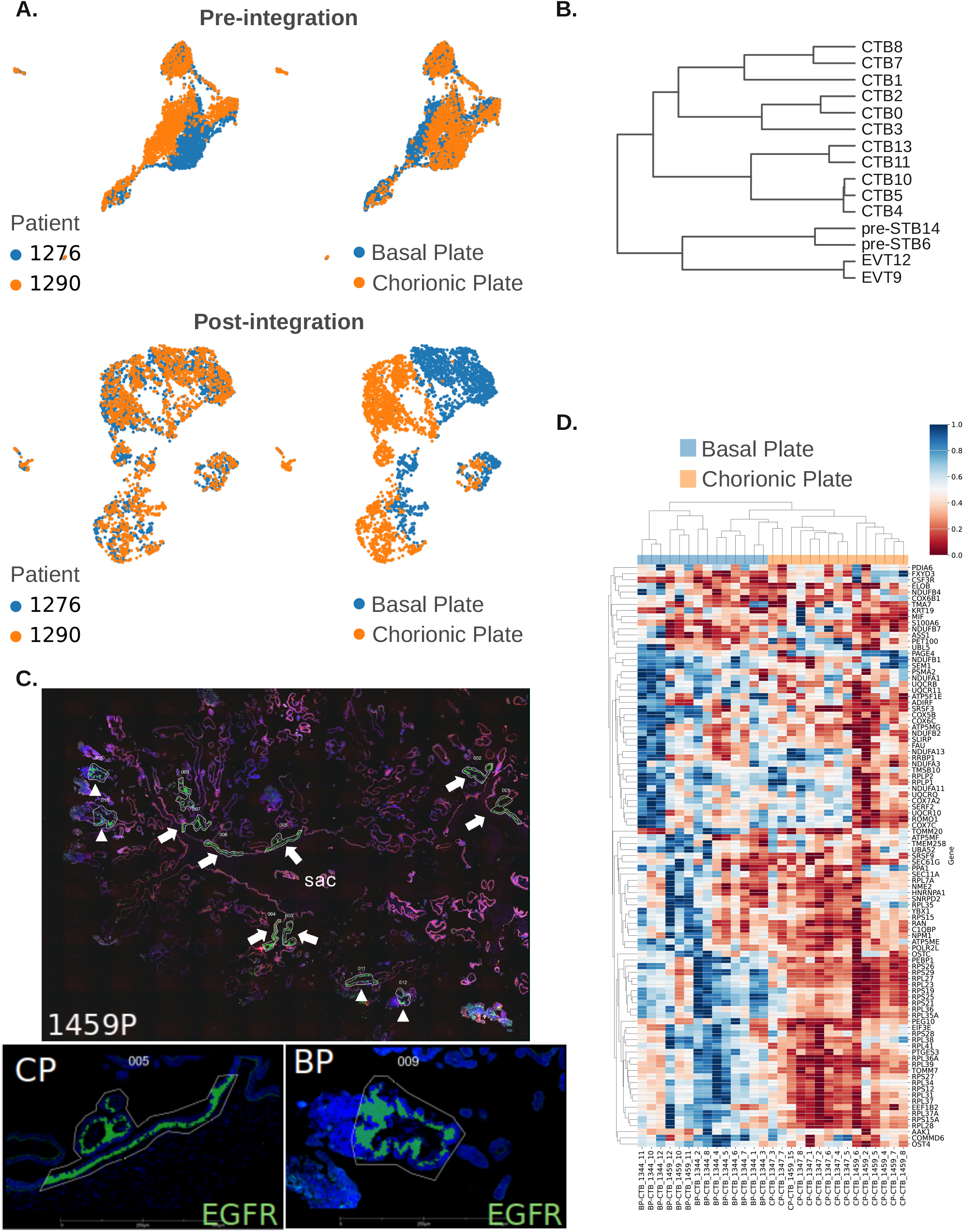
Basal and chorionic plate CTB. (A) UMAPs of the basal and chorionic plate samples before and after integration colored by the placenta number (left) and location of the sample (right). For each placenta, a basal and chorionic sample was sequenced. (B) Hierarchical clustering of annotated basal and chorionic plate clusters using all cells and genes post quality control filtering. (C) Representative scan of formalin-fixed paraffin-embedded placental tissue (placenta 1459P) used for GeoMx-based digital spatial profiling, showing tissue stained for EGFR to identify vCTB, with selection of Areas of Illumination (AOI’s) near the basal (BP, arrowheads) and chorionic plate (CP, arrows) based on spatial relation to the gestational sac. (D) Heatmap displaying the expression of genes that were determined to be differentially expressed between BP and CP CTB in 29 AOI’s, located within three different placentae, using the GeoMx whole transcription atlas (WTA) panel.

**Fig. S6.**
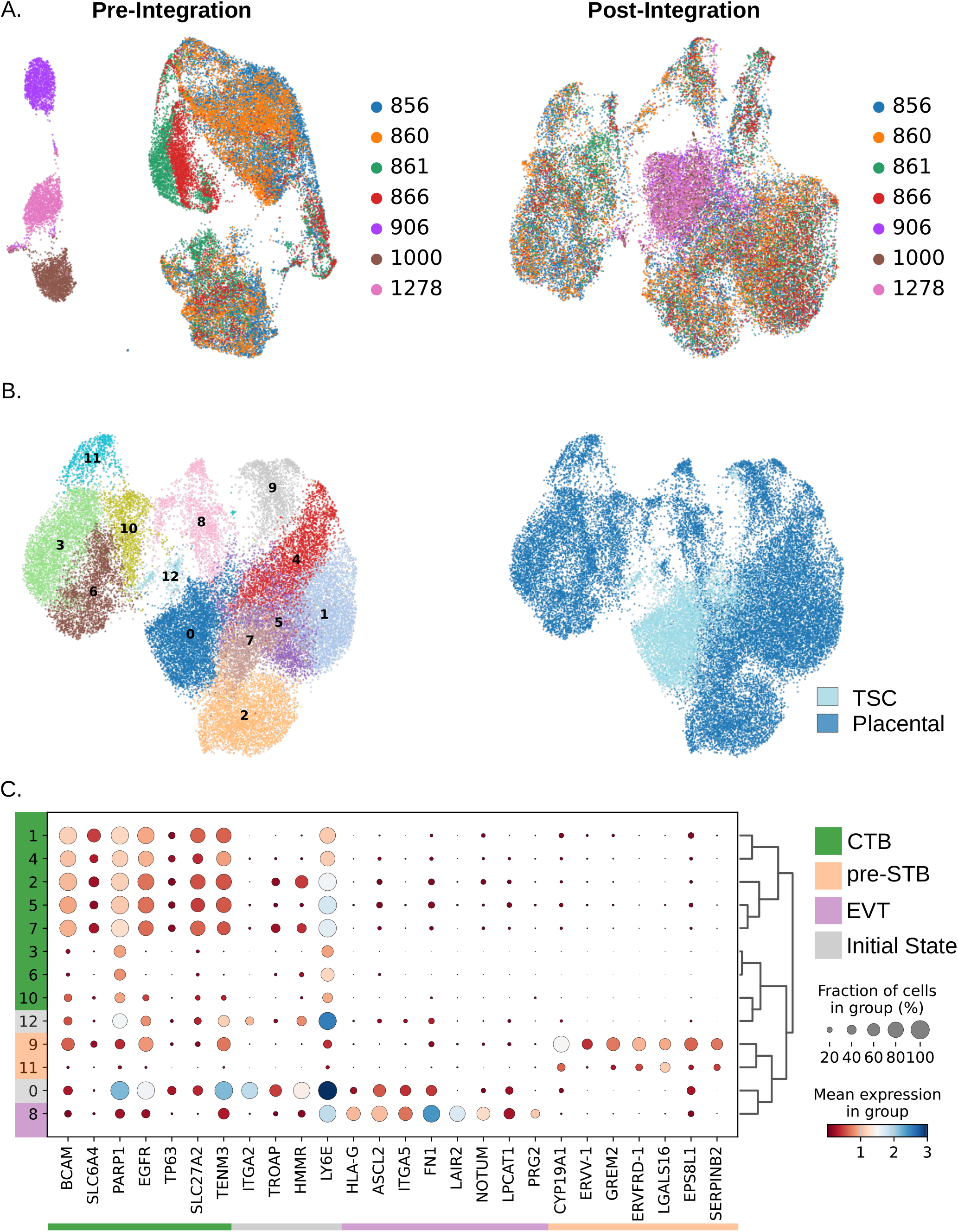
Trophoblast stem cell UMAPs. (A) UMAPs colored by the patient number, before (left) and after (right) integrating the placental and TSC samples together. (B) UMAPs post Leiden clustering, colored by cluster (left), vs. cell source (TSC or placental cell) (right). (C) Dot plot and hierarchical clustering of clusters using cell type-specific gene expression markers with CTB in green, EVT in purple, pre-STB in peach, and initial state markers in grey.

**Fig. S7.**
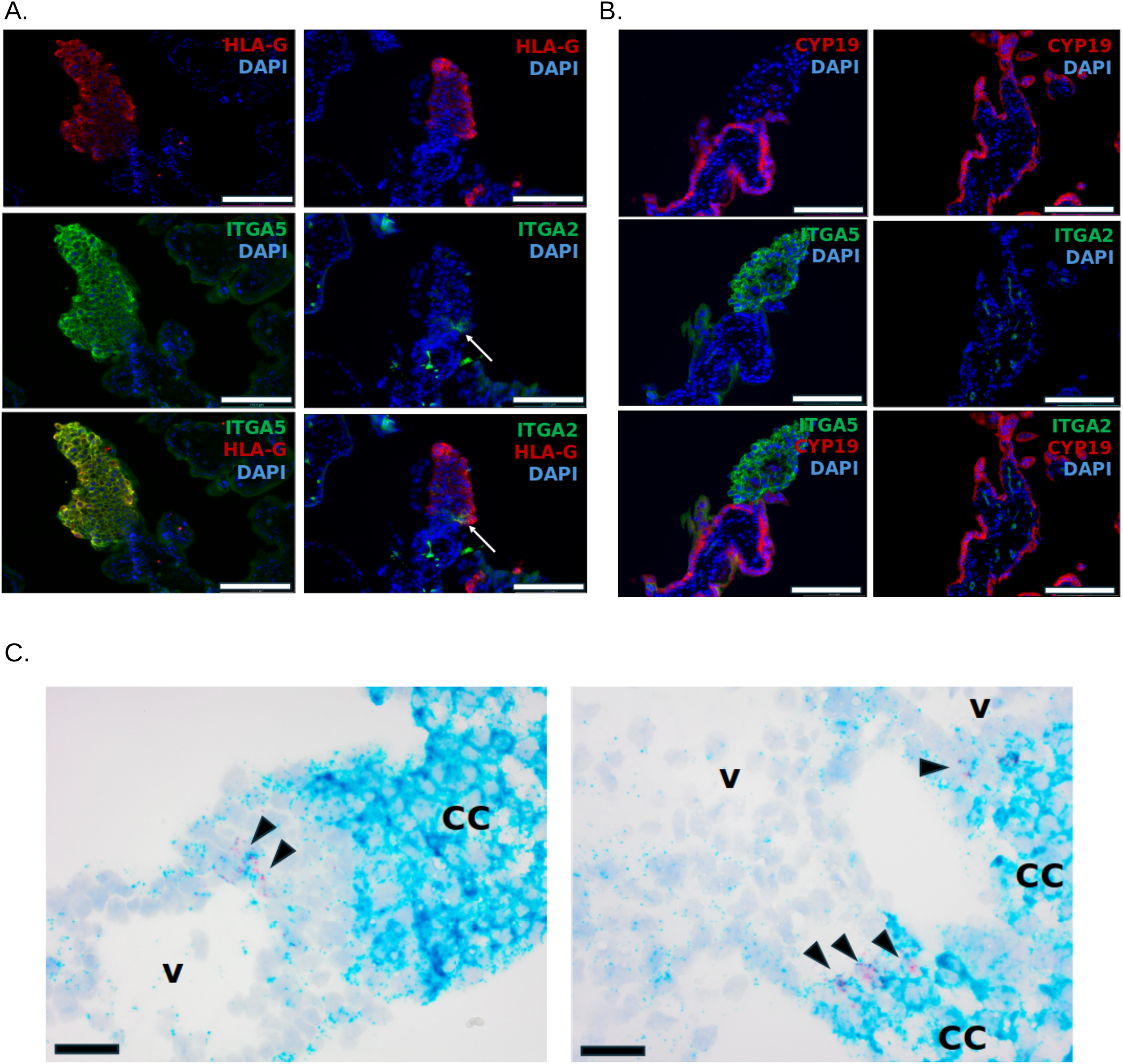
Confirmation of ITGA5 and ITGA2 expression in anchoring columns of first trimester CTB. (A) Immunohistochemistry of first trimester placenta with antibodies against ITGA5 and ITGA2, co-stained either with HLA-G (panel A) or CYP19A1 (B). ITGA5 is only expressed in column trophoblast, while ITGA2 is expressed in rare proximal column trophoblast (arrow in panel A), but also in fetal endothelial cells (panel B). Scale bar=156 *µ*m. (C) In-situ hybridization of first trimester placenta with probes against ITGA5 (teal) and ITGA2 (magenta). Rare dual-positive cells are noted in the proximal columns (arrowheads). V=villous core; CC=cell column. Scale bar=62.5 *µ*m.

**Fig. S8.**
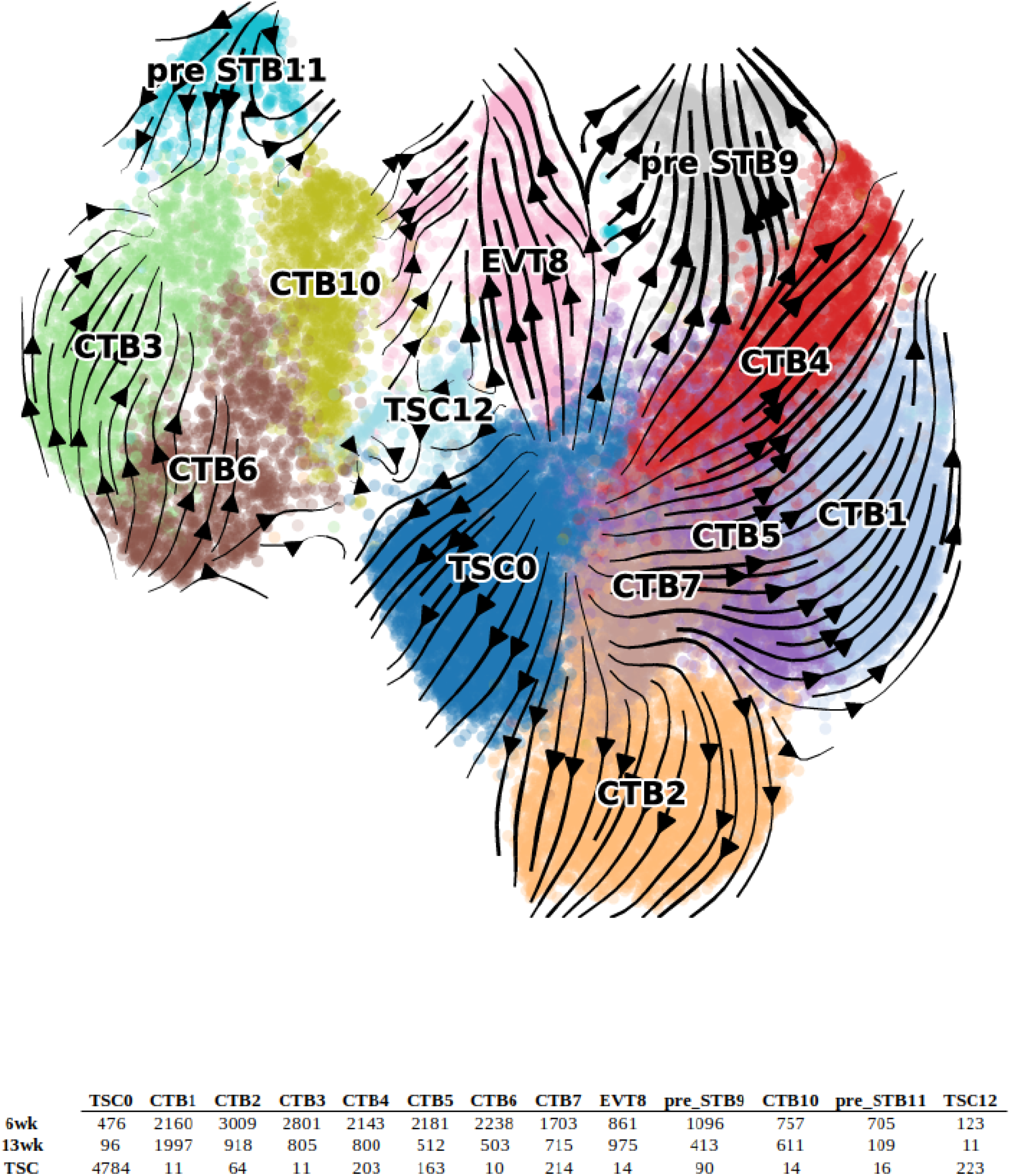
Trophoblast stem cell trajectory inference. RNA velocity projected as streamlines on the integrated UMAP. The UMAP contains both early and late first trimester CTB, as well as TSC’s.

**Fig. S9.**
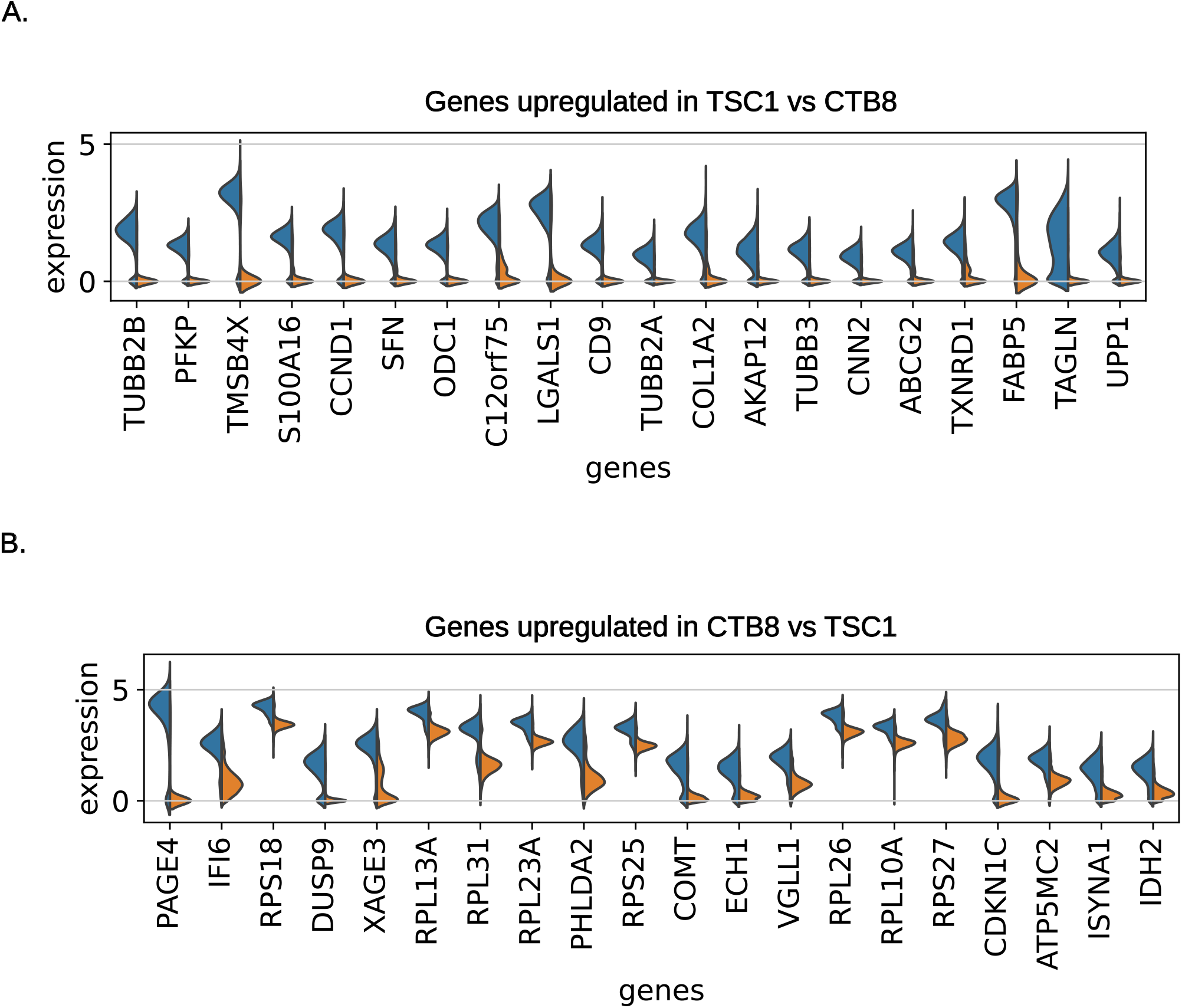
Differences in gene expression between TSC and CTB initial state clusters. (A) Violin plots of the top 20 upregulated genes in cluster TSC1 compared to CTB8. (B) Violin plots of the top 20 upregulated genes in cluster CTB8 compared to TSC1.

**Fig. S10.**
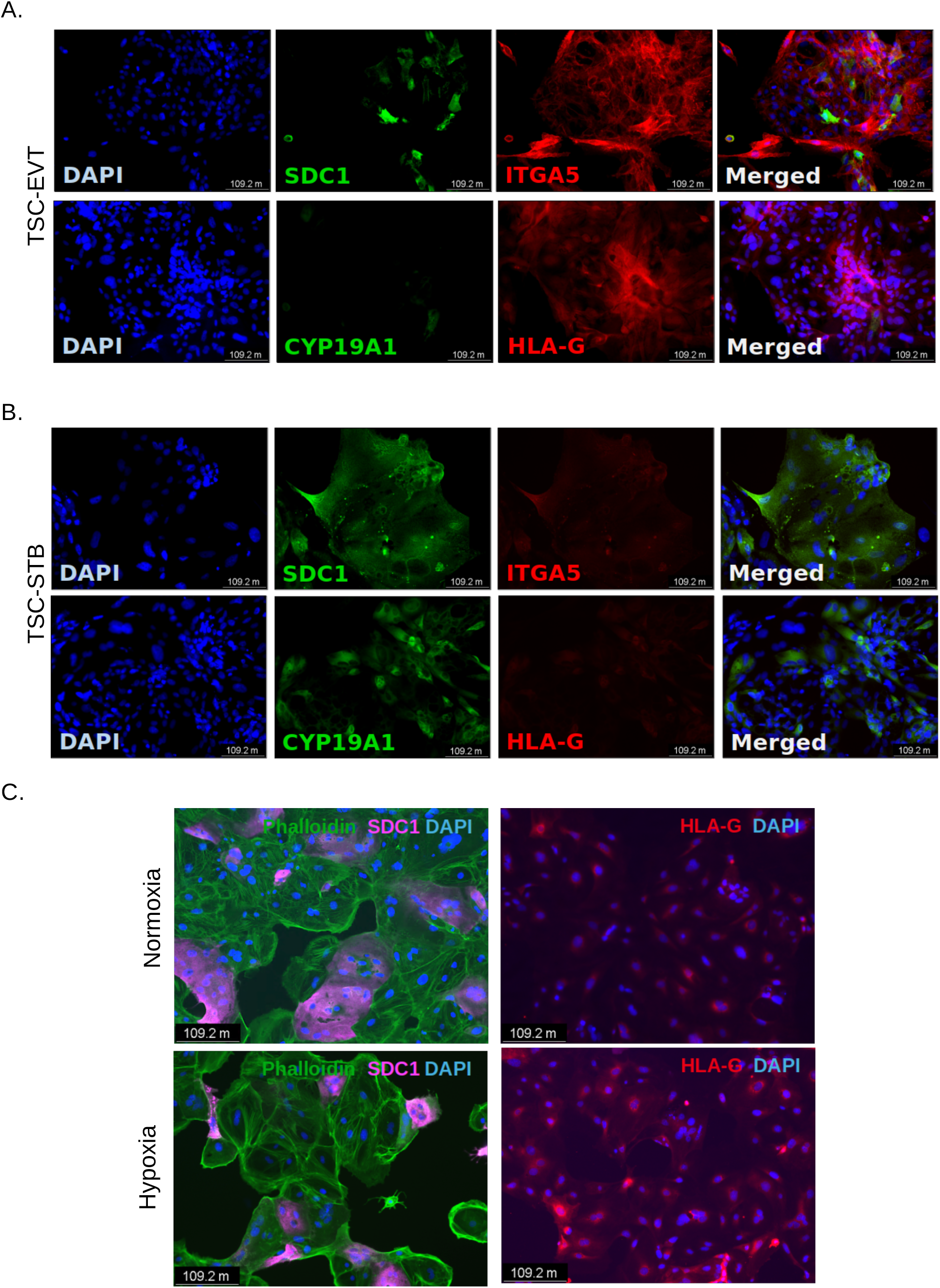
ITGA5 expression is lost when TSC are directed to differentiate towards STB. (A) Immunohistochemistry of STB markers (SDC1 and CYP19A1), pcEVT marker (ITGA5), and EVT marker (HLA-G) following differentiation of TSC to EVT. (B) Immunohistochemistry of STB markers (SDC1 and CYP19A1), pcEVT marker (ITGA5), and EVT marker (HLA-G) following differentiation of TSC to STB. (C) Immunohistochemistry of TSC showing large SDC1^+^ multinucleated cells when cultured in normoxia, and HLA-G^+^ mononuclear cells when cultured in hypoxia.

Table S7: Genes upregulated (> 1.5 Log2 fold change) in CTB3 (late) compared to CTB8 (early).

Table S8: Genes upregulated (> 1.5 Log2 fold change) in CTB10 (late) compared to CTB7 (late).

Table S9: Top 100 likelihood genes and the overlap between patients, directly upstream EVT and pre-STB clusters as determined by RNA velocity.

Table S10: Differentially expressed genes with a score over 15 upregulated in basal plate CTB compared to chorionic plate CTB.

Table S11: Differentially expressed genes with a score over 15 upregulated in chorionic plate CTB compared to basal plate CTB.

Table S12: Marker genes of all clusters after integration of primary trophoblast and TSC’s.

Table S13: Genes upregulated (> 1.5 Log2 fold change) in TSC1 vs CTB8.

Table S14: Genes upregulated (> 1.5 Log2 fold change) in CTB8 vs TSC1.

## Resource Availability

### Lead contact

Further information and requests for resources and reagents should be directed to, and will be fulfilled by, the lead contact, Mana Parast.

### Materials availability

This study did not generate new unique reagents. Requests for human trophoblast stem cell lines used in this study should be directed and will be fulfilled by the Lead contact, Mana Parast.

### Data and code availability

- Single-cell RNA-seq data and GeoMx digital spatial profiler data have been deposited at GEO and are publicly available as of the date of publication. Accession numbers are listed in the key resources table (GSE270174). Microscopy data reported in this paper will be shared by the lead contact upon request.
- This paper does not report original code. The code used for data processing and analysis has been deposited at Zenodo and is publicly available as of the date of publication. DOIs are listed in the key resources table.
- Any additional information required to reanalyze the data reported in this paper is available from the lead contact upon request.

## Experimental Model and Study Participant Details

### Patient recruitment and tissue collection

Human placental tissue samples for this study were collected under a UC San Diego Institutional Review Board-approved protocol; all patients gave informed consent for collection and use of these tissues. All samples were collected from pregnant individuals, undergoing elective termination of pregnancy, providing written informed consent. Early first trimester placental tissues, including those that were used to derive TSC and organoids, were taken from patients with gestational ages ranging from 5 weeks and 5 days to 8 weeks and 4 days. Late first trimester placental tissues (n=2) were taken from patients with gestational ages of 12 weeks and 2 days and 13 weeks (**Table S1**). Gestational age was determined based on crown-rump length, measured on ultrasound, and was stated in weeks/days from the first day of the last menstrual period. “Normal” sample was defined as a singleton pregnancy without any detectable fetal abnormalities on ultrasound.

The biological sex of the placental tissues used in this manuscript was not considered in the analyses presented here. The small sample size used in this study prohibited the authors from making meaningful inferences about biological sex differences. Sex determination can be inferred by those using the data in this study by expression of Y chromosome-linked genes.

### Trophoblast sample collection and preparation

Samples were processed within 2 hours of collection according to our established protocol(31). For whole placental samples, including TSC and organoid derivation, chorionic villi were minced and subjected to three sequential enzymatic digestions with Trypsin and DNase, followed by Percoll (Sigma-Aldrich) gradient separation. For single cell analysis, after filtration, cells were counted and loaded on the 10x Genomics chip. For basal and chorionic sample collection, we collected the villous tips (basal fraction) and the gestational sac (chorionic fraction) under a dissecting microscope, while the villi in the middle section were discarded. The two fractions were subjected to the same sequential enzymatic digestion and after filtration cells were separated on a Percoll (Sigma-Aldrich) gradient. Cell suspensions were loaded onto the 10x Genomics Chromium Controller for GEM generation. All single-cell libraries were constructed using the Chromium Next GEM Single Cell 3, Library & Gel Bead Kit v2 and v3 (10x Genomics), following the manufacturer’s protocol for all steps. Single-cell libraries were sequenced on an Illumina NovaSeq 6000 instrument. Capture rate and sequencing depth can be found in **Table S1**. scRNA-seq datasets are publicly available on the GEO repository (GSE270174). Leftover cells were lysed for RNA isolation using NucleoSpin kit (Macherey-Nagel) and proper basal vs. chorion separation was verified by enrichment of CDX2 (chorionic) and ASCL2 (basal) expression by qPCR. Libraries were prepared only from sample pairs showing correct marker enrichment.

## Method Details

### Patient-derived trophoblast stem cell establishment, culture, and differentiation to EVT and STB

Following Percoll (Sigma-Aldrich) separation, first trimester (GA 6-8 weeks; n=6) CTB were MACS-purified with either a PE-conjugated anti-ITGA6 antibody (Biolegend 313612) or an APC-conjugated anti-EGFR antibody (Biolegend 352906). MACS-purified CTB were then plated for TSC derivation as previously described(9, 13). Once cell morphology had stabilized (after 6-8 passages), TSCs were evaluated for expression of various surface markers, including EGFR, HLAG, and a panel of integrins by flow cytometry (see below).

Directed EVT differentiation was performed by plating 75,000 cells/well onto a 6-well plate pre-coated with 20ug/ml fibronectin in EVT differentiation media as described by Okae et al.(7). On day 3, EVT medium was replaced without NRG1, and Matrigel concentration was reduced to 0.5% until day 5. On day 5, cells were re-plated onto a 6-well plate pre-coated with 20ug/ml fibronectin using the latter media (without NRG1 and with reduced Matrigel). For immunostaining, cells were fixed on day 6.

Directed STB differentiation was performed by plating 25,000 cells/well onto a 6-well plate pre-coated with 2.5ug/ml collagen IV in the STB differentiation media as described by Okae et al.(7). Cells were fixed on day 6 for immunostaining.

For spontaneous differentiation, two TSC lines (1000P and 1048P) were plated at a density of 100,000 cells/well in a 12-well plate coated with 20ug/ml fibronectin and cultured in Dulbecco’s modified Eagle’s medium/F12 with 10% FBS under normoxia (21% O2) or hypoxia (2% O2, using an XVIVO X3 Workstation). After 4 days, cells were fixed for immunostaining, or collected for RNA isolation and qRT-PCR.

### Isolation and plating of first trimester cytotrophoblasts

CTB were isolated from the first trimester placental tissues using sequential digestion with Trypsin and DNase, followed by Percoll (Sigma-Aldrich) gradient separation as previously described(50). For the plating experiment, one million cells were plated per well in a 6-well plate, coated with 20ug/ml fibronectin, and cultured in Dulbecco’s modified Eagle’s medium/F12 with 10% FBS, 1x penicillin/streptomycin, and 50ug/ml gentamicin as previously described(50). Expression of a panel of integrins (ITGA6, ITGA5, and ITGA2) was evaluated by flow cytometry at the time of isolation (day 0) and after 4 days of culture in regular 37°C humidified incubator (21% oxygen) with 5% CO2.

In a separate set of experiments, CTB were sorted for ITGA5 by magnetic-activated cell sorting (MACS), following the manufacturer’s protocol (Miltenyi). ITGA5 positive and negative cells were then plated onto 20ug/ml fibronectin-coated plates and cultured again in DMEM/F12 with 10% FBS and antibiotics under normoxia (21% O2) or hypoxia (2% O2, using an XVIVO X3 Workstation). Cells were then fixed for immunostaining.

### Trophoblast organoid derivation and culture

CTBs were isolated from first trimester placental tissues using sequential digestion with Trypsin and DNase, followed by Percoll (Sigma-Aldrich) gradient separation as previously described(50). Isolated trophoblast cells were sorted for ITGA5 by MACS with ITGA5-FITC antibody (Biolegend 328008), following the manufacturer’s protocol (Miltenyi). ITGA5 positive and negative cells were counted and plated into Phenol Red free Matrigel (Corning) domes at 37,000 cells/15ul domes in trophoblast organoid media (TOM) comprised of Advanced DMEM/F12 (Life Technologies), 1x N2 and 1x B27 supplements (Life Technologies), 2mM Glutamax (Life Technologies), 1.25mM N-Acetyl-L-cysteine (Sigma), 500nM A83-01 (Tocris), 1.5µM CHIR99021 (Sigma-Aldrich), 80ng/mL human R-spondin 1 (Peprotech), 50ng/mL human EGF (RD), 50ng/mL human HGF (Stemcell Technologies), 2.5µM prostaglandin E2 (PGE2) (Selleck Chemicals), 100ng/mL human FGF2 (BioPioneer), and 2µM Y-27632 (Selleck)(15). Cultures were maintained in a 37°C humidified incubator with 5% CO2 and media changed every 2-3 days. After 10-12 days, organoids were passaged following mechanical dissociation as described in Turco et al.(15). After the first passage, organoids were maintained in TOM media and passaged with a combination of TryPLE Express (Thermo Scietific) treatment for 3-5 min. followed by mechanical dissociation every 10-14 days and re-seeded at 20,000 cells/20µL in Matrigel dome(51).

### Flow cytometry analysis

Isolated CTB and derived TSCs were stained, either individually with EGFR-APC (Biolgend 352906), HLAG-PE (ExBIO 1P-292-C100), ITGA6-AF647 (Biolegend 313610), ITGA5-FITC (Biolegend 328008) and ITGA2-AF594 (RD FAB1233T), or using a multi-color panel of EGFR-PE/Cy7 (Biolegend 352910), HLAG-APC (ExBIO 1A-292-C100), ITGA1-PE (Biolegend 328304), ITGA2-AF700 (RD FAB1233N), ITGA5-FITC (Biolegend 328008) and ITGA6-BV421 (Biolegend 313624), with relative isotype control antibodies, for 1 hour in flow cytometer wash buffer (1% FBS, 2%BSA and 0.03% Sodium Azide in PBS) in the dark. After staining, cells were washed three times with wash buffer and fixed with 2% PFA, and samples were acquired on either a FortessaX20 or Fortessa (BDLSRFortessa) flow cytometer (BD Biosciences). Data analysis was performed using FlowJo and data were represented as bar graphs using GraphPad Prism.

### Immunofluorescent staining of cells and tissues

Cells were fixed with 4% paraformaldehyde (PFA) in PBS for 10 minutes, washed once with PBS, then permeabilized with 0.1%Triton X for 10 mins at room temperature (RT), and washed three times with PBS. Cells were blocked with 5% Goat serum and 0.05% Triton X in PBS for 1 hour at RT. Cells were stained overnight at 4°C with primary antibodies, including mouse anti-SDC1 (0.5ug/ml, Abcam ab34164), rabbit anti-ITGA5 (1.1ug/ml, Abcam, ab150361), and mouse anti-CYP19A1 (1.0ug/ml, Novus Biologicals, NBP3-07826), and Rabbit anti-HLAG (0.5ug/ml, Abcam ab283260), followed by secondary antibodies (Alexa Fluor-conjugated goat anti-rabbit or anti-mouse antibodies, Thermo Fisher Scientific) for 1 hour at RT in the dark. Nuclei were stained with DAPI, and images were taken on a Leica fluorescence microscope.

Immunofluorescent staining was also performed on formalin-fixed paraffin-embedded (FFPE) placental tissues. 5-mm serial sections were cut, dried, and deparaffinized and rehydrated using xylene and ethanol. Sections were permeabilized in a 0.1% triton solution in 1x TBS for 10 minutes, then subjected to antigen retrieval in EDTA at 100°C for 15 minutes. Tissue sections were blocked using 10% normal goat serum (Jackson ImmunoResearch, Cat 005-000-001) and 5% BSA (Gemini BioProducts, Cat700-110) in TBST for 30 minutes at 37°C. Serial sections were stained with mouse anti-HLAG (1.0ug/mL, Abcam, ab283270), rabbit anti-ITGA2 (1.0ug/mL, Bioss, bsm-52613R) and rabbit anti-ITGA5 (1.1ug/ml, Abcam, ab150361) and incubated at 4°C overnight. The following day, slides were washed with 1x TBS then incubated with Alexa Fluor 488 and 594 conjugated goat anti-rabbit and goat anti-mouse (Thermo Fisher) for 1 hour at room temperature covered from light. Slides were then washed in a 1:5000 dilution of DAPI in 1x TBS for 7 minutes, washed in 1x TBS, then mounted using Prolong Gold (Invitrogen). Images were acquired using a Leica fluorescence microscope and individual channel photos were merged using ImageJ.

### *In situ* hybridization

First trimester placental tissue samples were fixed in a neutral-buffered formalin and embedded in paraffin wax. Five µm sections were subjected to in-situ hybridization on a Ventana Discovery Ultra automated stainer (Ventana Medical Systems) at the UC San Diego Advanced Tissue Technology Core laboratory. RNAscope probes specific to human PAGE4, HMMR, TROAP, LY6E, ITGA5, and ITGA2 were purchased from ACD-Bio. Following amplification steps, the probes were visualized using an HRP-based reaction, visualizing single probes (PAGE4, HMMR, TROAP, and LY6E) in brown and dual probes in teal (ITGA5) and red (ITGA2); slides were counterstained with hematoxylin. Slides were visualized by conventional light microscopy on an Olympus BX43 microscope (Olympus).

### GeoMx digital spatial profiler whole transcriptome assay

For spatial transcriptomic analysis, three 6-week gestation placentas, previously formalin-fixed and paraffin-embedded, were selected from the Center for Perinatal Discovery’s biorepository (**Table S1**). Samples containing a visible, relatively intact embryonic sac were selected so that the basal and chorionic areas were easily identifiable. 5µm sections were mounted on SuperFrostTM Plus slides and used within 2 weeks. Sections were deparaffinized and incubated overnight with the Whole Transcriptomic Assay UV-cleavable biological probes according to manufacturer’s instructions. Samples were stained with CONFIRM® anti-EGFR (5B7) Rabbit Monoclonal Primary Antibody (Ventana 790-4347, 1:20 dilution) and anti-HLA-G (4H48) mouse primary antibody (Abcam ab52455, 1:1000 dilution) for 2 hours followed by secondary staining for 1h with Alexa Fluor anti-rabbit-647 (Life Technologies, 1:500 dilution), Alexa Fluor antimouse-594 (Life Technologies, 1:500 dilution), and SYTO13 (Nanostring, 1:10 dilution) for nuclei staining. Slides were scanned on a GeoMx digital spatial profiler (Nanostring). Based on the morphology markers, areas of interest (AOIs) containing at least 100 EGFR^+^/HLA-G^-^ nuclei were manually selected. In total, we identified 35 AOIs, 16 from the basal plate and 19 from the chorionic plate. Samples were collected and libraries prepared according to the manufacturer’s instructions. Libraries were pair-end sequenced on a NovaSeq 6000 in the UCSD IGM core at 3 million reads per sample. Fastq files were converted into DCC files following the GeoMx NGS pipeline (v.2.0.0.16). Data QC was performed on the GeoMx DSP software according to standard parameters (minimum 100 nuclei per AOI, area size at 1000µm2, negative probe count to 1, sequencing saturation at 30, Q3 normalization) and then exported for further downstream analysis. Genes validated with the GeoMx data were differentially expressed in either the basal or the chorionic clusters with a log2 fold change over 0.5 and a score over 15 using Scanpy’s rank-genes-groups method with default parameters (**Supplementary Table S10 & S11**). Data are publicly available on the GEO repository (GSE270174).

### RNA isolation and qRT-PCR

On day 4, cells were collected, and RNA isolated using NucleoSpin® (Macherey-Nagel, USA) kit. 300ng of RNA was reverse transcribed to prepare cDNA using Prime-Script™ RT reagent kit (TAKARA, USA) following the manufacturer’s instructions. qRT-PCR was performed using Power SYBR® Green RT-PCR Reagents Kit (Applied Biosystems, USA) using the following primer pairs in a 5’ to 3’ orientation: ASCL2 forward CACTGCTGGCAAACGGAGAC, ASCL2 reverse AAAACTCCAGATAGTGGGGGC; CGB forward ACCCTGGCTGTGGAGAAGG, CGB reverse ATGGACTCGAAGCGCACA; ITGA1 forward CTGGACATAGTCATAGTGCTGGA, ITGA1 reverse ACCTGTGTCTGTTTAGGACCA; L19 forward AAAACAAGCGGATTCTCATGGA, L19 reverse TGCGTGCTTCCTTGGTCTTAG. Data were normalized to L19 and shown as fold-change over undifferentiated TSC. GraphPad Prism was used to perform the statistical analysis. Ordinary One-way ANOVA with multiple comparison correction were used to calculate the statistical analysis after comparing with undifferentiated TSC. Data is expressed as mean ± standard deviation of 2-ddCt values. The level of statistical significance was set at p < 0.05.

### Trophoblast scRNA-seq library construction and sequencing

Cells were run on the 10X Genomics platform with the Chromium Next GEM Single Cell 3’ kit. Samples and libraries were prepared following the manufacturer’s protocol to attain 5,000 to 10,000 cells per reaction. Libraries were pooled and sequenced using the Illumina NovaSeq 6000 sequencer. Capture rate and sequencing depth can be found in **Table S1**.

### Single-cell RNA-seq data analysis

#### A. Data mapping and technical artifact removal

All scRNA-seq data was mapped using the STARsolo function of STAR (v.2.7.10a)(52). STARsolo performed error correction and demultiplexing of cell barcodes using user-input whitelist (3Mfebruary-2018.txt for version 3 of the 10X Genomics Chromium System and 737K-august-2016.txt for version 2 data), mapped reads to the reference genome (GRCh38, version 32, Ensembl 98), deduplicated and error corrected UMIs, quantified per-cell gene expression, and spliced and unspliced reads similar to Velocyto. The following are the CellRanger mimicking commands used: –outFilterScoreMin 30 –readFilesCommand zcat –soloCBmatchWLtype 1MM_multi_Nbase_pseudocounts –soloUMIfiltering MultiGeneUMI_CR –soloUMIdedup 1MM_CR – soloCellFilter EmptyDrops_CR –soloUMIlen 12 –soloType CB_UMI_Simple –soloCBwhitelist 3M-february-2018.txt –clipAdapterType CellRanger4 –outFileNamePrefix name –outSAMattributes All–outSAMtype BAM SortedByCoordinate –quantMode GeneCounts –soloCBlen 16 –soloCBstart 1 – soloUMIstart 17 –soloFeatures Gene GeneFull SJ Velocyto. Following STARsolo quantification, the data was run through CellBender’s (v. 0.3.0) ‘remove-background’ command(19) to remove systematic biases and background noise due to ambient RNA molecules and random barcode swapping.

#### B. Data pre-processing, filtering, integration, and quality control

The data was then loaded into Scanpy (v. 1.10.1)(22) for preprocessing, filtering, and quality control. Doublet removal was performed by Scrublet(20) with an expected doublet rate of 0.076. Following doublet filtering, cells with less than 200 genes expressed were removed and genes detected in less than 3 cells were removed. Next, cells with greater than 20% mitochondrial DNA content were removed as well as cells with greater than 5% ribosomal gene expression or 15% hemoglobin gene expression. The data was then normalized and the cell cycle was regressed out using the cell cycle genes from Macosko et al.(53), and highly variable genes were selected. The data was then integrated using scvi-tools (v. 1.1.2)(21), specifically using the scVI (single-cell Variational Inference) model with “batch” and “cell_source” as the categorical covariate keys. Integration was performed before clustering but after combining all datasets being analyzed and performing the above quality control steps.

#### C. Clustering and clustering annotation

Integrated datasets were clustered using Scanpy’s Leiden clustering algorithm, an improved version of the Louvain algorithm(54), with a resolution of 0.75 unless otherwise specified. Marker genes were calculated using default parameters and clusters were annotated as trophoblast if they expressed all or some of the genes *KRT7, GATA3, PAGE4, TFAP2A, HLA-G*, or *CYP19A1* and lacked all or most of the expression of the genes *HLA-A, HLA-B, HLA-DRA, CD14, VIM*, or *CD34*. The non-trophoblast clusters were then removed from downstream analyses and the remaining trophoblast clusters were reclustered using the same resolution after the nearest neighbors distance was recalculated and embedded using UMAP. Trophoblast clusters were annotated using the following gene expression: CTB (*BCAM, SLC6A4, PARP1, EGFR, TP63, SLC27A2*, and *TENM3*); EVT (*HLA-G, ASCL2, ITGA5, FN1, LAIR2, NOTUM, LPCAT1, PRG2*, and *AOC1*); Pre-STB (*CYP19A1, ERVV-1, GREM2, ERVFRD-1, LGALS16, EPS8L1*, and *SERPINB2*). One cluster shared expression of both EVT and pre-STB markers and was therefore defined as STB-EVT-like. Hierarchical clustering using Pearson correlation method with complete linkage was used to assess the relative distance between clusters. Differential expression between two groups was performed using ‘rank_genes_groups’ and setting one cluster as the reference and the other as the ‘group’. Genes were considered up- or downregulated using a log2 fold change of 1.5 unless otherwise stated.

#### D. Trajectory inference and initial state determination

Trajectory inference was performed using scVelo (v. 0.3.2), an RNA velocity(55) analysis toolkit for single cells that leverages splicing kinetics to recover directed dynamic information using an expectation-maximization framework(23) or a deep generative model(56). A function from https://github.com/JBreunig was used to put together all the spliced and unspliced data before preprocessing and moments calculations were performed. Velocities were estimated in a gene-specific manner using the stochastic mode for all trajectories except for the individual patient trajectories, which used the dynamical mode.

To compute the initial state, we used CellRank2 (v. 2.0.4)(24), a Markov state modeling framework for cellular dynamics, and the velocity kernel, which computes a transition matrix based on RNA velocity, along with the Generalized Perron Cluster Cluster Analysis algorithm (GPCCA estimator)(25). When computing the initial state for the dataset containing the combined early- and late placental samples, first the velocity kernel and connectivity kernel were combined in an 0.8/0.2 ratio. Then using the GPCCA estimator, the terminal states were set to the EVT and pre-STB clusters and then the number of states was selected to be four. From these four states, the initial state was predicted. For the basal and chorionic dataset, the GPCCA estimator and the velocity kernel were used along with a Schur decomposition to find 8 macrostates, one of which was determined to be the initial state. A similar process was used with the TSC and early integrated dataset, but with the initial state being chosen from three macrostates.

### Quantification and Statistical Analysis

Data are reported as mean ± standard deviation. Statistical analyses were performed using GraphPad Prism. Ordinary one-way ANOVA and multiple comparison corrections were used to calculate the statistical significance compared to undifferentiated TSC. qPCR data were analyzed following the ddCt method: data were normalized on the housekeeping gene L19 and expressed as fold change (2^-ddCt^) over undifferentiated TSC. The level of statistical significance was set at p < 0.05. For scRNA-seq statistical analyses, please refer to the applicable section in the methods details.

